# Mera: A scalable high throughput automated micro-physiological system

**DOI:** 10.1101/2022.08.30.505827

**Authors:** Finola E. Cliffe, Conor Madden, Patrick Costello, Shane Devitt, Sumir Ramesh Mukkunda, Bhairavi Bengaluru Keshava, Howard O. Fearnhead, Aiste Vitkauskaite, Mashid H. Dehkordi, Walter Chingwaru, Milosz Przyjalgowski, Natalia Rebrova, Mark Lyons

**Affiliations:** Hooke Bio Ltd, L4A Smithstown Industrial Estate, Shannon, Co. Clare, Ireland, V14 XH92; Pharmacology and Therapeutics, Biomedical Sciences, Dangan, NUI Galway, Galway, Ireland; Centre for Advanced Photonics and Process Analysis, Munster Technological University, Cork, Ireland, T12 P928

**Author notes:** E-mail address: ^a^.

**Keywords:** High throughput screening, Microfluidics, Preclinical Drug Screening, Microphysiological Systems

## Abstract

There is an urgent need for scalable Microphysiological Systems (MPS’s)^1^ that can better predict drug efficacy and toxicity at the preclinical screening stage. Here we present Mera, an automated, modular and scalable system for culturing and assaying microtissues with interconnected fluidics, inbuilt environmental control and automated image capture. The system presented has multiple possible fluidics modes. Of these the primary mode is designed so that cells may be matured into a desired microtissue type and in the secondary mode the fluid flow can be re-orientated to create a recirculating circuit composed of inter-connected channels to allow drugging or staining. We present data demonstrating the prototype system Mera using an Acetaminophen/HepG2 liver microtissue toxicity assay with Calcein AM and Ethidium Homodimer (EtHD1) viability assays. We demonstrate the functionality of the automated image capture system. The prototype microtissue culture plate wells are laid out in a 3 × 3 or 4 × 10 grid format with viability and toxicity assays demonstrated in both formats. In this paper we set the groundwork for the Mera system as a viable option for scalable microtissue culture and assay development.

## 1 Introduction

Despite the complex and costly preclinical drug screening process only 10% of these candidate molecules will made it through clinical trials and to the market (Van Norman, 2019b) highlighting the need for more accurate models in this field. The existing biochemical and cellular assays play an important role in refining the list of potential drug candidates from thousands to tens by enabling researchers to focus on biochemical pathways and whole cell level metrics. However, these assays give little insight into the complex and dynamic interactions that are at play in a whole organism. Animal models on the other hand were thought to offer a more complex environment to study efficacy and toxicity of drug candidates and as such have become a mandated part of the drug testing pathway. Multiple studies have shown that animal models are poor at predicting potential toxicity and incapable of predicting efficacy (Olson et al., 2000, Tamaki et al., 2013, Van Norman, 2019a). Genetic diversity, gene expression profiles, post translational modification and inter-individual diversity are not captured in ‘clonal’ animal models

Over the last 15 years cells culture models have become increasingly sophisticated. Advances in 2D cell culture have come in the form of micro-patterned coculture (Berger et al., 2015, March et al., 2015). This technique is compatible with 96 well SBS plate formats and involves the specific spatial location of different cell types relative to one another. Successful examples include HepaRG hepatocytes with 3T3 murine embryonic fibroblasts showing sensitivity and specificity results above those of primary hepatocytes in a tier 1 compound screen (Ware et al., 2021).

Interestingly, Microphysiological Systems (MPS’s) have increasingly shown their relevance as models for drug toxicity and efficacy screening (Hofer and Lutolf, 2021, Kim et al., 2020, Shankaran et al., 2021). Such models have the potential to enable researchers to make better informed decisions earlier in the drug development pipeline possibly before utilising animal models thereby implementing the 3R’s strategy to reduce, refine and possibly replace these models in the future.

MPSs of nearly every major human organ including hosting liver (Asif et al., 2021, Chen et al., 2021, Ewart et al., 2022, Görgens et al., 2021, Takebe et al., 2013), lung (Sachs et al., 2019, Ainslie et al., 2019), intestine (Sato et al., 2009, Peters et al., 2020, Jalili-Firoozinezhad et al., 2019), neural (Lancaster et al., 2013, Liu et al., 2019, Pamies et al., 2017) and cardiac tissue (Rogozhnikov et al., 2016, Conant et al., 2017, Lind et al., 2017) (amongst many others) have been developed using cells from a variety of sources including cell lines, primary cells and stem cells and have been grown using single or multiple cell types. The complexity that brings power and versatility of these systems comes with the challenge of having to define, mature and measure outcomes in a meaningful and well controlled manner. These systems generally have been cultured in traditional microplate formats which are scalable and have many existing technologies to support them from robotic plate handlers and pipetting systems to plate readers and automated optics systems.

However, while there are a number of arguments for performing MPS experiments in the traditional formats researchers interested in whole organism interactions with drug candidates may require a more sophisticated approach. Organ to organ communication via chemical mediators is a ubiquitous phenomenon in the human body. For example, hepatokine expression from the liver including sex-hormone-binding-globulin (Ding et al., 2009) and hepassocin (Li et al., 2010) correlate with type 2 diabetes and liver regeneration respectively. Muscle expressed myokines such as IL-6 have been shown to have impacts on liver, adipose and endocrine tissues (Pedersen and Febbraio, 2008). Vernetti et. al. (2017) investigated the organ processing of three compounds, terfenadine, trimethylamine-N-oxide and vitamin D3 by various interconnected organ models. They demonstrated the organ specific processing of these compounds was consistent with clinical data. When new drugs are investigated it may be the intention of the researcher to supply the drug in a pro-drug format so that its metabolised form is active to enable targeted drug delivery or conversely the drug may be metabolised by one or more organs such as the liver and the metabolites go on to have toxic effects on other tissues (Edington et al., 2018, Herland et al., 2020).

To model these effects *in vitro* a more sophisticated flow path than exists in a single well may be desirable. Organising organoids in individual wells connected by a common fluidic channel with the capacity to drive flow across the tissues would go a long way to recapitulating the structure and mechanical arrangement as seen in the human body. Many microfluidic organ-on-a-chip and body-on-a-chip solutions have been demonstrated in labs across the world (Ahadian et al., 2018, Ma et al., 2021, Picollet-D’hahan et al., 2021). Scaling these technologies to make them widely available can be challenging from a manufacturing point of view though there have been some notable successes such as Emulate’s Zoe culture model and organ chips (Emulate Bio, 2022) and TissUse’s Humimic Starter device, AutoLab and associated chips (TissUse GmbH, 2022). Choice of materials is also under scrutiny with challenges around small molecule absorption in materials such as PDMS being flagged as challenges for the future of the MPS industry.

Even in the later stages of pre-clinical drug screening drug developers are in the position of needing to screen tens if not hundreds of drug candidates and scaling MPS technologies while maintaining the physiological advantages they offer can be challenging. The balance of number of tests a system provides inevitably pulls against the complexity the system can offer. To address this challenge we developed a two mode system (though other modes are possible) based around the Mera lid configuration. By using different lid designs, we sought to answer the question of how a system can have fluid pathways that allow a common tissue type to be matured and then the pathways reorientated so that the fluid pathway creates a circuit across tissue types as would be seen in a body-on-a-chip model. To facilitate operation at scale, automation of environmental control, fluidics and optics are key features enabling these technologies. The Mera system includes integrated electronic valving and fluid pumping controllable through a graphical user interface, a custom designed Bioplate which is SBS compatible, automated environmental control and automated optics for image capture. We demonstrate the functionality of these systems using HepG2 liver spheroids which will be herewithin referred to as microtissues (MTs) and a paracetamol toxicity assay in both a 3×3 and 4×10 well Bioplate prototypes.

## 2 Materials and Methods

### 2.1 Materials

10% Fetal bovine serum (FBS), Dulbecco’s modified Eagles medium (DMEM), 1% Penicillin/Streptomycin, L-Glutamine, Phosphate buffered Saline (PBS), Trypsin-EDTA, Acetaminophen, Ethidium Homodimer (EtHD1) and DMSO were purchased from Sigma-Aldrich (Dublin, Ireland). Calcein AM viability stain was obtained from BD Biosciences (Berkshire, UK).

### 2.2 Cell culture and microtissue formation

HepG2 human hepatocellular carcinoma cells were obtained from ATCC (ATCC-HB-8065, LCG Standards, Teddington, UK) and were maintained in DMEM supplemented with 10% FBS and 1% Penicillin/Streptomycin. Cells and microtissues were cultured at 37°C in a humidified, 5% CO_2_ atmosphere and were passaged upon reaching 70-80% confluency.

HepG2 cells were plated on PrimeSurface 96-well ultra-low attachment plates (PHC Europe B.V., Loughborough, UK) at a density of 200 cells/well, centrifuged for 5 min at 200 x g and incubated in 5% CO_2_ at 37°C for microtissue formation. Microtissues formed within 24 hours and culture medium was refreshed every 2-3 days by removing 100uL of medium and replacing with fresh media until approx. day 7 when transfer to the Mera Bioplate occurred (Figure 1, A). All microtissues were formed in ULA plates and cultured to approx. 220um diameter (approx. 7 days) prior to experimentation.

**Figure 1:**
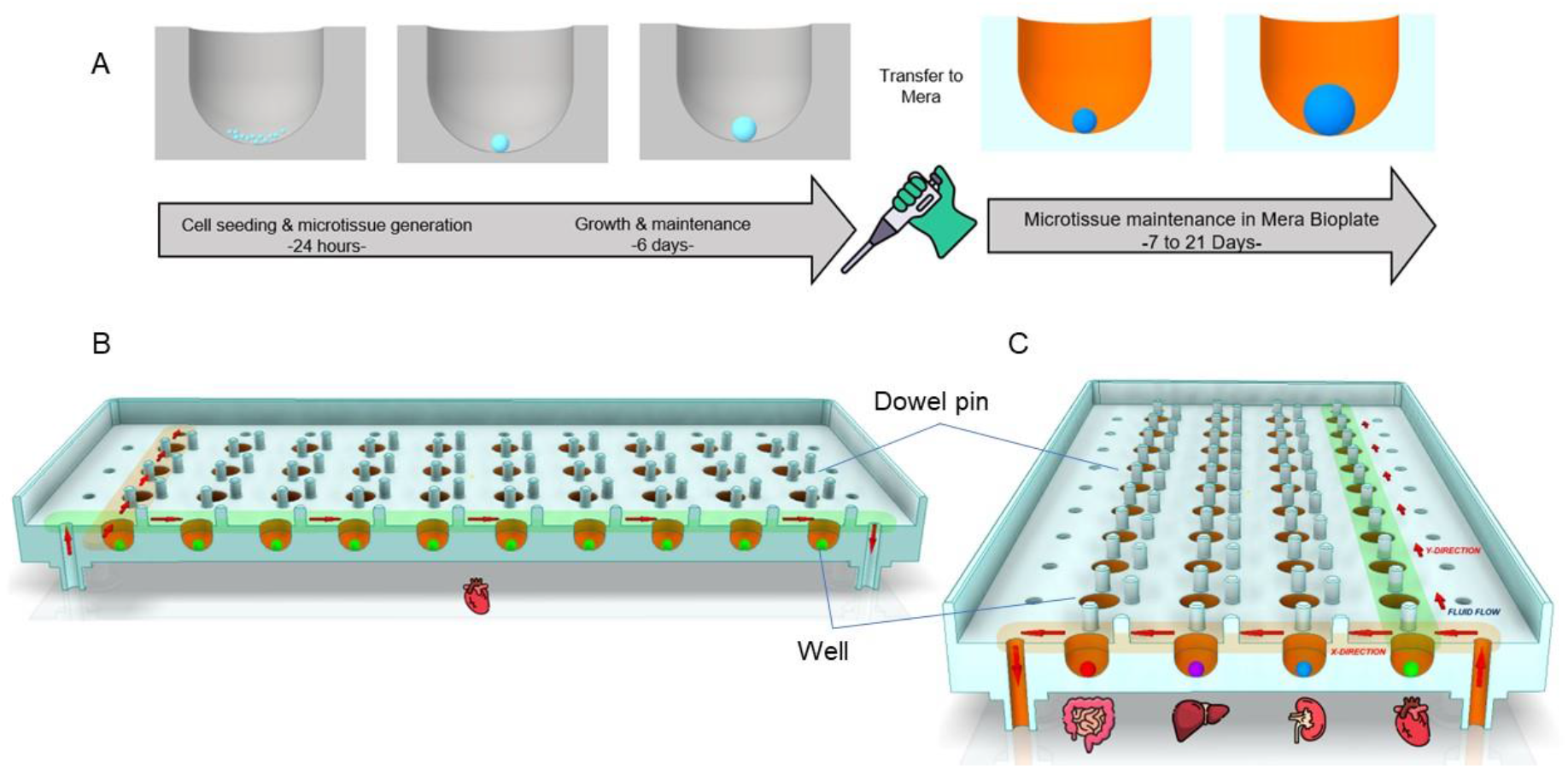
A) Microtissue preparation and transfer to Bioplate timeline, (B) Layout of the Bioplate 4 × 10 grid design detailing media flow for microtissue growth, C) Layout of the Bioplate 4 × 10 grid design showing media flow direction for intertissue communication via recirculation perfusion. Organ images reproduced with permission from Flaticon.com.

### 2.3 Description and Device Fabrication/Assembly

The organs of interest in the human body can be modelled by specific microtissues while microfluidic perfusion is used to model fluid circulation in the human body. The concept of the MPS is illustrated in Figure 1, B & C with an example of a four-organ system on a 40 well plate termed the Bioplate. In this paper the authors focus on demonstrating a scaled back version of this concept by assessing a single organ HepG2 liver model within a perfused MPS. The system comprises two main aspects, the Bioplate, which is essentially a culture plate, and the Mera system, which controls the perfusion through the Bioplate via a series of pumps and valves.

The fluidic assembly of Mera consists of four main parts, i) the Lid, of which there are two different versions, the 40 well Bioplate (4 well x 10 channel format & 10 well x 4 channel format, Figure 2B) and one version of the 9 well Bioplate (3 wells x 3 channels), ii) The flexible membrane, iii) the Bioplate which is arranged in a rectangular format consisting of either a 10 × 4 or 3 × 3 grid containing round bottomed wells into which microtissues can be situated and iv) the baseplate. Each well on the Bioplate has two neighbouring dowel pins (See Figure 1 and Supplementary information Figures 2 & 3). To assemble the fluidic system a sandwich is made using the lid, the membrane, the Bioplate and the baseplate (Figure 2A & D). The dowel pins act like poles in a marquee to hold the membrane in position above the floor of the Bioplate creating a channel through which fluid can flow. The ridged walls of the lid compress the membrane against the edges of the Bioplate forming channels in X and Y formats (Figure 2B) depending on which lid layout has been employed (4 well x 10 channel, 10 well x 4 channel or 3 well x 3 channel format). The inlets and outlets of each given channel are found along the periphery of the Bioplate and form the entry and exit points for the fluid to run through the channels. Up to 40 microtissues can be cultured on the 4 × 10 well Bioplate while nine microtissues can be accommodated on the smaller 3 × 3 well Bioplate.

**Figure 2:**
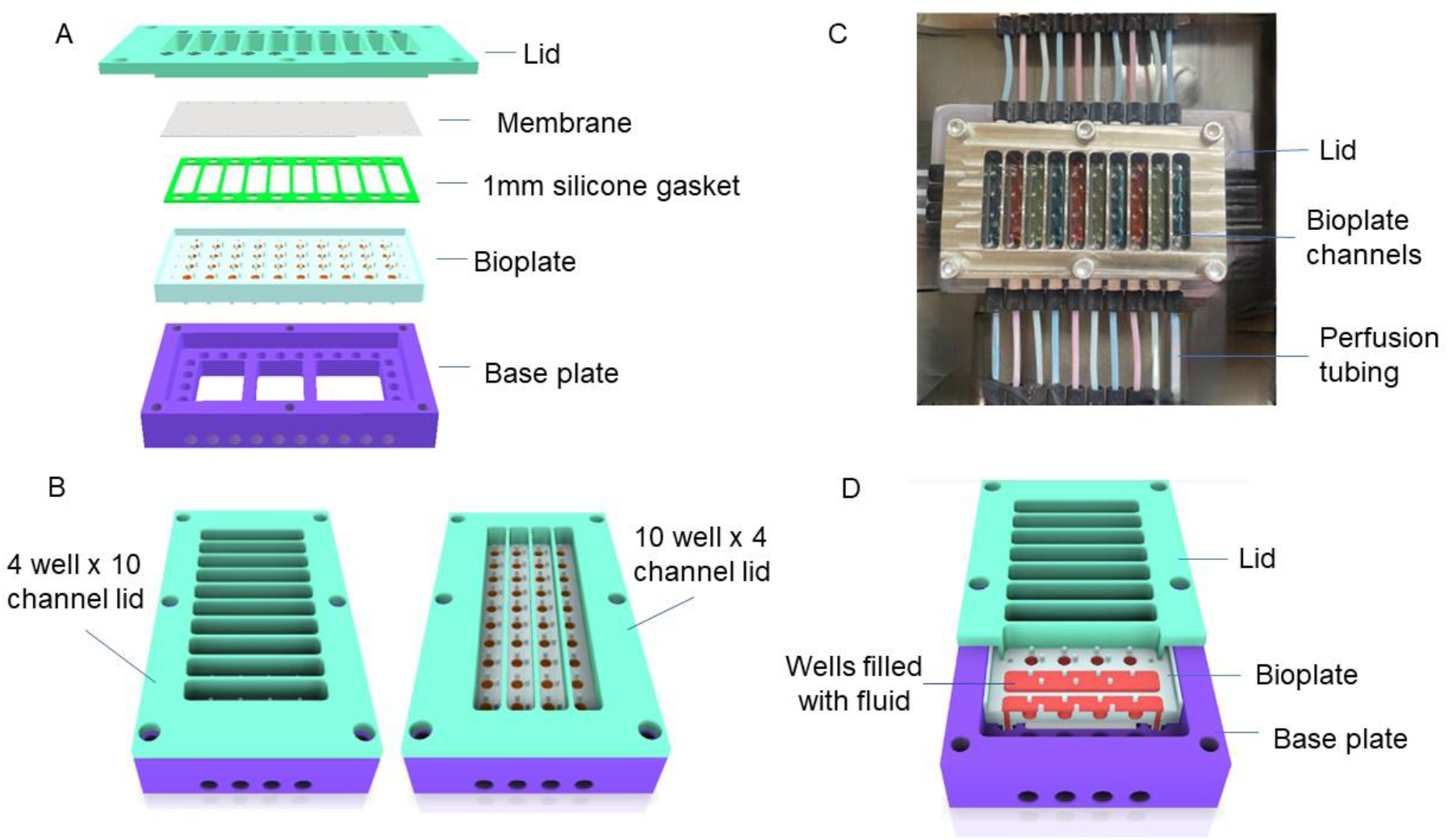
Bioplate and assembly, A) Illustration of Bioplate assembly. The Bioplate is inserted into the baseplate, and the lid is placed onto the assembly along with a membrane and silicone gasket to create channels for fluid perfusion, B) Illustration of Bioplate with optional lid formats which are employed to change the direction of fluid flow, 4 wells x 10 channels and 10 wells x 4 channels, C) Image of Bioplate with channels alternately filled with red and blue food dye integrated with baseplate and lid, D) Cutaway illustration of core Mera fluidic system with integrated Bioplate, baseplate and lid (10 rows x 4 wells).

### 2.4 Machining

All manufactured components were fabricated in-house using a Hermle 5-axis milling machine (Hermle C250, Gosheim, Germany). All parts were designed using AutoCAD Inventor software (ProCAD Engineering, Limerick, Ireland) prior to conversion into a machining program complete with tooling and G-codes (Hypermill, OPEN MIND). All materials in contact with cells within the system were tested for biocompatibility by incubating HepG2 cells for up to 7 days in the presence of the material of interest and cell morphology and viability was subsequently assessed (Supplementary information Figure 4 & 5).

### 2.5 Bioplate Design and Fabrication

The Bioplate (PMMA, Engineering Steels Limerick, Ireland) was designed in two formats, the 3×3 (9 wells) and 4×10 (40 wells) grid versions. Each Bioplate consists of wells entirely independent of each other where fluid is supplied via an automated peristaltic perfusion system comprising of a fixed number of inlets specific to the plate design, inlet/outlet tubing and connections and collection reservoirs. The Bioplate consists of either 9 (3 × 3 grid format) or 40 (4 × 10 grid format) U-bottomed wells. The design of the Bioplate module has several advantages, 1) the dimensions and concave shape of the wells (2mm radii hemispheres) localises the microtissues to the centre of the well to facilitate more accurate imaging, consistent nutrient supply and decreasing the possibility of damage due to shearing forces 2) it also allows direct access for efficient loading of microtissues into the wells, manipulation during experiments or removal for downstream analysis 3) dead volume can be kept to a minimum 4) the optical transparency of the base of the wells enables high-quality optical readouts. All well compartments have a diameter of 4mm, a height of 3.5mm and a volume of 35uL (excluding overlying perfusion channel volume). The perfusion channels for 4 × 10 (4 wells) are 7mm wide, approx. 1mm high and 44.6mm long and for the 10x 4 (10 wells) are 7mm wide, approx. 1mm high and 100mm long. A single channel (consisting of four wells) has a combined volume of approx. 312uL while the perpendicular channel (consisting of 10 wells) has a combined approx. of 700uL. The perfusion channels for 3 × 3 are 7mm wide, approx. 1mm high and 35mm long with a combined volume of approx.245uL. The height is an average measurement as the membrane is not a uniform height throughout the channel due to the position of the dowel pins.

### 2.6 Bioplate integration with Mera

Mera consists of a central base plate and associated lid (and sealing gaskets) with optional recirculation functions into which the Bioplate sits. Perfusion to the central plate is supplied by a valve block manifold with associated fluidic fittings and perfusion control valves. This is all supported on a stainless-steel support structure.

The Bioplate is integrated into the base plate with the lid to create a complete assembly (Figure 2). The Base plate (17-4 Stainless Steel, Engineering Steels Limerick, Ireland) houses the Bioplate, whilst facilitating the connections to the liquid supply. The lid formats are made from stainless steel (17-4 Stainless Steel, Engineering Steels Limerick, Ireland). Both are designed with open channels to facilitate fluid flow while also providing surface area to accommodate gaseous diffusion. Of the lid variations, one can be employed to facilitate fluid flow across 10 wells in a four-channel format (10 wells x 4 channels i.e. hosting microtissues of the same organ type) and the second for facilitating flow across four wells in a 10-channel format (4 wells x 10 channels, i.e. hosting microtissues of different organ types). The 3 well x 3 channel format lid can be employed in both directions to facilitate flow as it is a square format. These lids provide the structure to create a physical barrier between adjacent channels, preventing any fluidic or material crossover. A membrane (adhesive layer of breath easy polyurethane (PU) membrane from Diversified Biotech, Dedham, MA, USA or a 1mm silicone membrane, (60 shore from TYM Seals & Gaskets, Devizes, UK) can be attached to the underside of the lid; this allows gaseous exchange to occur via osmosis whilst constructing a channel within which fluid can pass. When employing the PU membrane, the physical barrier of the membrane is further strengthened using a 1mm thick silicone gasket (60 shore, TYM Seals & Gaskets, Devizes, UK) which also alleviates the impact of the compressive force of the steel lid onto the acrylic insert, decreasing the risk of damage to the insert. The polyurethane membrane is single use however all other components including the silicone membrane and gasket (for use with PU membrane) can be sterilised and reused. The lid is attached to the base plate using six M6 threaded hex socket bolts.

Fluid enters the base plate from the valve block manifolds via ¼ -28” threaded IDEX fittings (CIP-XP-301X, Darwin Microfluidics, Paris, France). The fluid is directed 90 degrees upward through the internal geometry of the base plate, which facilitates the delivery of fluid to the Bioplate. The fluidic seal between the base and the Bioplate is maintained with O -rings to tolerance as per the radial compression design recommendations (Parker, 2015).

### 2.7 Perfusion control valves

The 4.5mm width of the thirty-two 24V 2/2 Flipper-Solenoid Valves (Type 6650, Burkert, Ingelfingen, Germany) allowed the channels to be placed closer together than larger alternatives, whilst also providing the basis for the Ø1.4mm internal fluidic channels (max diameter tolerated by the valve specifications). Please see https://www.burkert.com/en/type/6650 for operational details. The thirty-two valves comprised of two entry valves, two exit valves and twenty-eight inner directional valves, which facilitated the control of the direction of fluid movement.

### 2.8 Valve block manifolds

Each valve block (Acrylic, Engineering Steels, Limerick, Ireland) can hold two Burkert valves; there is a male and female block to complete the valve block manifold assembly, into which ¼ -28” IDEX fittings (CIL-XP-301X, Darwin Microfluidics, Paris, France) are connected. The fluidic seal between the individual blocks within the assembly is maintained with O-rings to tolerance as per the radial compression design recommendations. As an added precaution, an M3 threaded rod intersects the entire valve block manifold. An M3 nut on the distal ends of the rod provides a compressive force to complement the fluidic sealing capacity of the O-rings.

To minimise the introduction of air into the system, an inlet/outlet T-block (PMMA, Engineering Steels, Limerick, Ireland) was designed to omit tubing from the design whilst maintaining the functionality of a directional switch valve. Three ¼ -28” IDEX fittings (CIL-XP-301X, Darwin Microfluidics) are threaded into the block to complete the connection to the system.

The entire system is housed on a block of Aluminium (Engineering Steels Limerick, Ireland). This allows parts to be fastened in place, providing a neat solution and consistency during experiments by reducing moving parts. The assembly of the Mera system is depicted in Figure 3.

**Figure 3:**
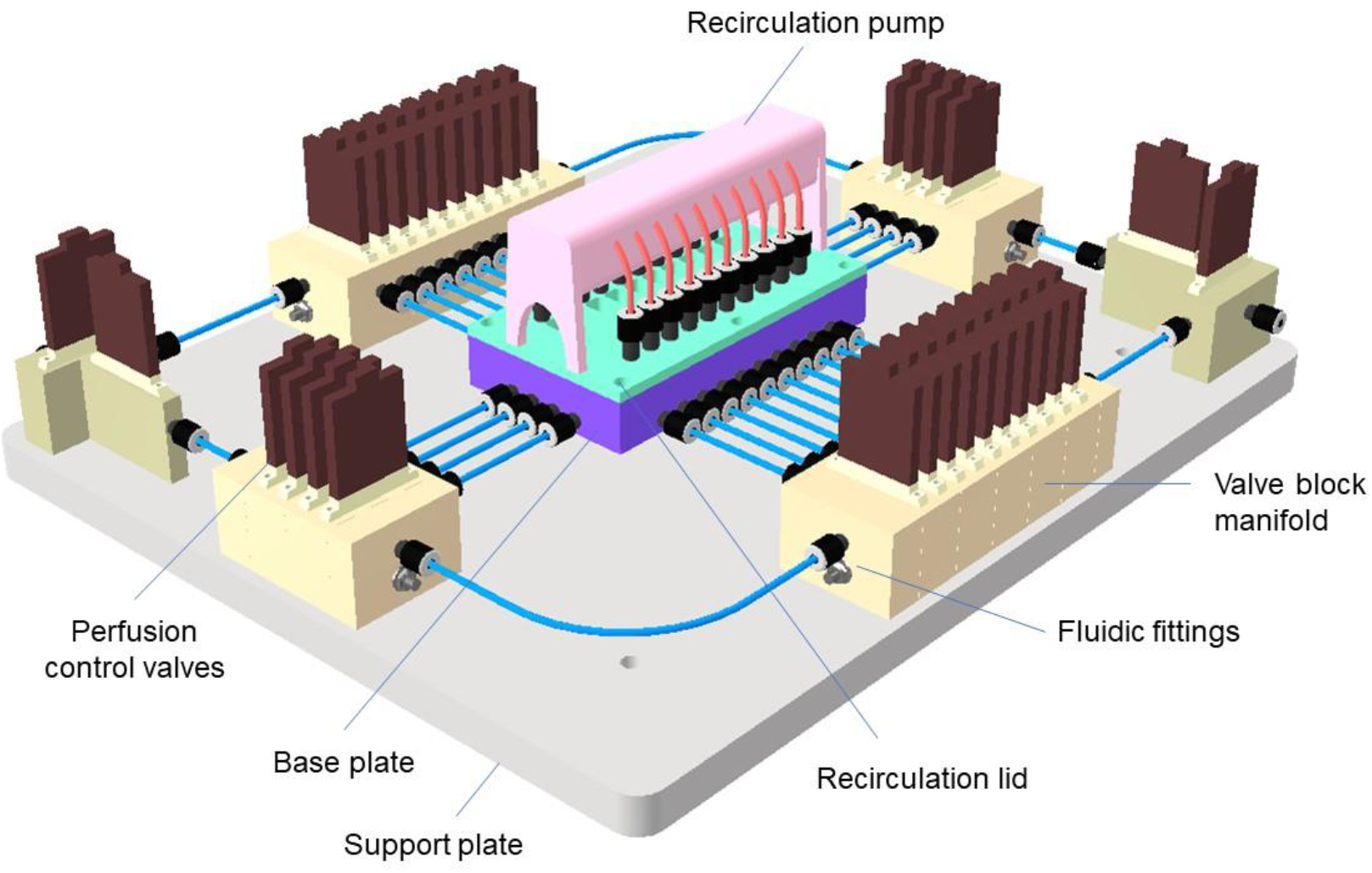
The Mera System with associated components identified.

### 2.9 Perfusion control

Two peristaltic pumps (KCM-B168, Kamoer Fluid Tech, Shanghai, China) are used to control perfusion through the system (Supplementary Figure 1). A constant rate of approximately 2.5ml/min was used to prime the system while a slightly lower flow rate of 2.0mL/min was used to feed, drug and stain the microtissues perfused across each line for a minute once per hour. This rate can be adapted depending on the individual needs of the cell model. A third pump (Takasago Electric Inc, Nagoya, Japan) used for recirculating media is in the validation stage. The recirculation line is approximately 300mm long. The recirculation pump runs at the same rate as the peristaltic pumps, ensuring a consistent environment for MT’s. The system is being tested to replicate current results while diminishing the requirement for media from an external source. The fluidic lines consist of PTFE tubing (1/8” OD, 1/16” ID) (BL-PTFE-3216-20, Darwin Microfluidics, Paris, France).

### 2.10 Imaging system

A custom epi-florescence microscope was designed in-house and incorporated within the environmental chamber for automated *in-situ* imaging of microtissues. The microscope setup is depicted in Figure 4A & B. The optical assembly of the imaging module consists of a 470nm blue colour (M470L5, Thorlabs, Germany) and a 530nm green colour (M530L4, Thorlabs, Germany), an LED light source for excitation of the sample, a filter cube (excitation filter, emission filter and dichroic mirror), a camera tube (WFA100, Thorlabs, Germany) and an objective lens with 5x magnification (46143, Mitutoyo, Japan). A CMOS camera (acA1920-155ucMED, Basler AG, Germany) was used for microscopy imaging. The microscope was integrated with an automated 3-axis motion control system. A python-based imaging GUI (Figure 4B) was developed using a Basler Pylon library to acquire the images and automatically control the motion of the microscope. This microscope was used online to monitoring of MT growth and placement within the wells of the Bioplate.

**Figure 4:**
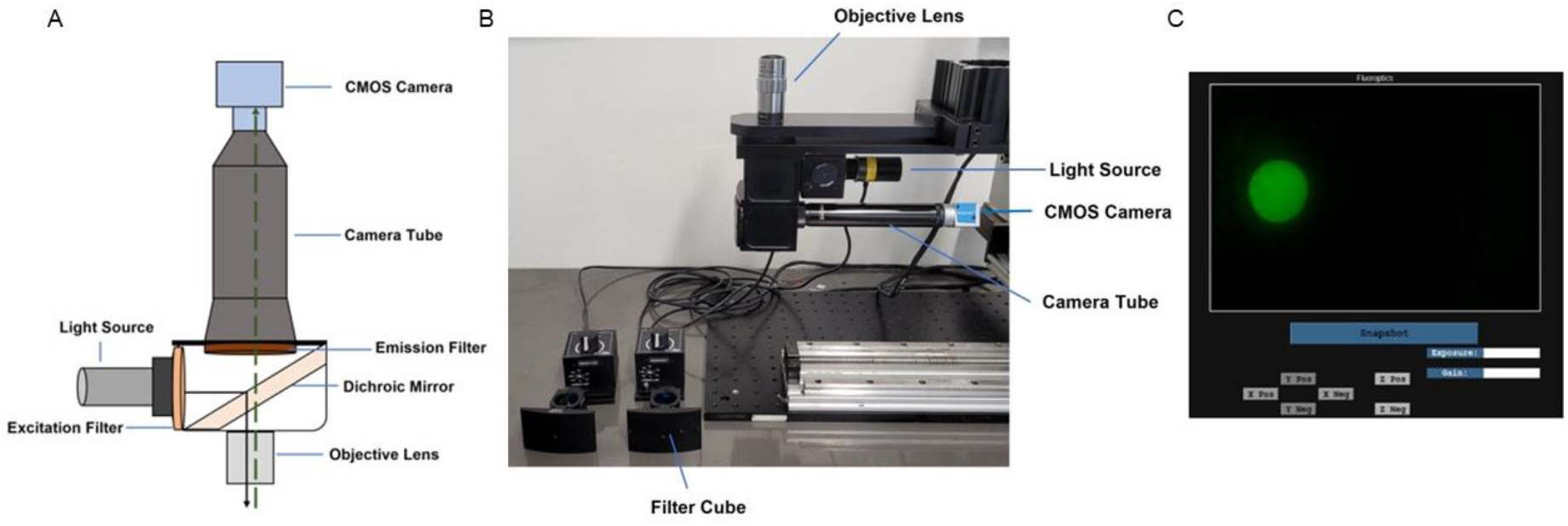
A) Epi-fluorescence microscope assembly illustration, B) Image of epi-fluorescence microscope assembly, C) Python-based graphical user interface (GUI)

For reproducibility, brightfield and fluorescence images were acquired for all microtissue data using an IX70 Olympus inverted microscope (Olympus, Tokyo, Japan) with a 4x objective and an AmScope Microscope Digital Camera (MU1003, AmScope, UK). All images were analysed using the Z-stacker App (CAPPA-CIT, Cork, Ireland) which allows for intensity-based image analysis for user selectable regions of interest. It was developed using OpenCV (Open-source Computer Vision library), a library of programming functions aimed mainly at machine vision and image analysis applications and consists of the following process, image alignment, sharpness map calculation, map comparison and image composition.

### 2.11 Environmental Chamber

To grow and maintain reproducible microtissues, mammalian cells require a temperature of 37°C and 5% CO_2_ within a sterile environment. To achieve this, an environmental chamber with a dimension of 1.5m x 1m x 2m was fabricated using stainless steel (SQ Fabs. Limerick, Ireland). The chamber was designed to be easily accessible for Bioplate removal, maintenance, and sterilization procedures. The Mera system and imaging system integrated within the environmental chamber is depicted in Figure 5A & B. A temperature control system consisting of a digital temperature sensor (DS18B20, DFRobot, China) and compact enclosure heater fans (02801.1-01, STEGO, Germany) were integrated to maintain a temperature of 37°C while a 12V DC axial fan was mounted on top of the enclosure heater fans to increase airflow velocity. A laminar flow system consisting of a HEPA filter and a centrifugal fan is built into the environmental chamber.

**Figure 5:**
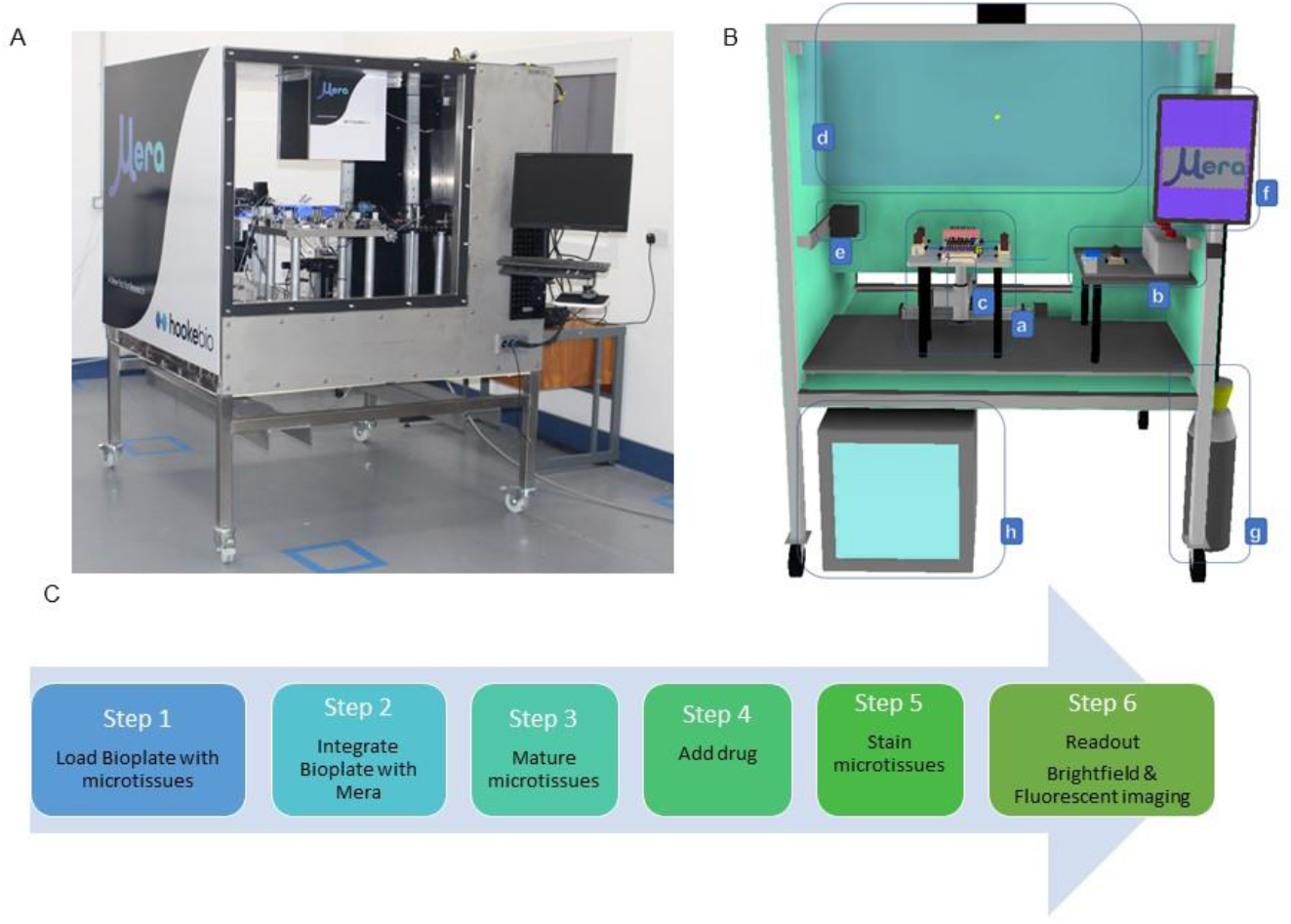
A) Image of the Environmental chamber, B) Illustration of the Environmental chamber integrated with Mera, imaging system and associated components; a) Mera system, b) fluidic and perfusion control system, c) imaging system, d) laminar flow unit, e) heater fans f) python-based GUI, g) CO_2_ supply, h) reagent refrigeration unit, C) Mera system steps involved in the experimentation process.

CO_2_ was supplied in a 6kg VB canister (BOC, Limerick, Ireland) which was safety tethered adjacent to the unit. CO_2_ sensor output is required to control the activation of the CO_2_ system. A high concentration, low power CO_2_ sensor (2091-EXPLORIR-WHV-E-100-ND, Gas sensing Solutions Ltd, UK) and a two-port solenoid valve (VX230AG, SMC corporation,) are used to control the inflow of the CO_2_ gas to the environmental chamber.

Reagents including media and buffers were stored in a miniature fridge which was retrofitted to include a length of tubing to supply fluids to the system, the length of tubing is elongated to include sufficient slack to ensure the fluid rises to 37°C via radiant heat within the hour timeframe between doses.

### 2.12 Software and Electronics

A microcontroller board based on ATmega2560 (Arduino mega2560. Arduino, Italy) was used as a processing unit to control temperature and CO_2_ levels in the environmental chamber and peristaltic pump and solenoid valves of the fluidic subsystem of the Mera system.

A python-based graphical user interface (GUI) was developed to control the environmental chambers environmental conditions, including the fluidic system’s perfusion rates and automated 3-axis motion control of the epi-fluorescence microscope.

### 2.13 Mera operation and testing

The Bioplate, silicone gasket and lid were cleaned using 70% IPA and sterilised using UV for 15 min before MT seeding and loading. The Mera system was sterilised by perfusing 70% IPA through the system for 20 minutes, followed by PBS washing prior to commencing the experiment. Liquid disinfection using IPA is employed as it is not only cost-effective but ensures decontamination of internal tubing and valves. The system is compatible with non-heat based commercially available sterilisation procedures such as chemical sterilisation e.g. ethylene oxide (EtO) and irradiation methods such as gamma ray or electron beam. Heat sterilisation using dry heat or steam should be avoided due to potential damage to sensors. The experimentation steps are outlined in Figure 5C.

HepG2 MTs were loaded into the Bioplate by carefully transferring them from the ULA plate using a single-channel pipette. The Bioplate was then inserted into the base plate, ensuring the O-ring seals were engaged at each of the connections. The system was primed using an automated multi-stage programme. Using this priming programme, all bubbles were removed before commencing the experiment. This began by removing any potential bubbles in the supply line by directing approximately 5mL of fluid around the external circuit of the system until the fluid reaches the far side of the manifold block. Fluid was then flushed through each of the Bürkert valves individually until the Bioplate was filled. This added a volume of approx. 24mL sitting in the channel above the wells, displacing any bubbles in the wells to the top of the channel where they burst. The silicone layer was submerged into this fluid onto the face of the Bioplate acting as a gasket seal. The channels were sealed with an adhesive polyurethane membrane (Diversified Biotech, Dedham, MA, USA) or silicone sheet attached to the base of the lid, which was aligned with the top silicone gasket and the lid was then fastened to the base plate. Any excess fluid was pushed out of the system via tubing into a disposable collection unit, ensuring no bubbles remain in the sealed system. This membrane allowed the exchange of oxygen, carbon dioxide and water vapour; prevents contamination, evaporation and fluid leakage and allows normal cell growth to proceed.

Nutrient medium (serum-free) or media containing 20mM Acetaminophen (APAP) was supplied to the MTs through the inlets and perfused at an average flow rate of 2.0 mL/min ± 0.02 (n=10) per line for 72h. After 72 hours, PBS buffer was perfused through the system to replace the media and subsequently, the MTs were stained by perfusing multiplexed 5μM calcein AM/10μM EtHD1 stain and held stationary for 3 hours. A corresponding control was undertaken on a 96 well microplate which was manually stained with 3μM Calcein AM and 10μM EtHD1 and held for 3hr in the incubator. Calcein stain concentrations varied due to differences in the experimental format and setup which can affect diffusion characteristics. Even staining of calcein AM and EtHD1 was observed across all wells in the Bioplate thereby confirming that diffusion occurs within the wells as uptake of the stain only occurs when it is in close proximity to the cell.

The MTs were washed with PBS twice to remove traces of media prior to the addition of stain. At the end of the time period, the Bioplate was removed from the system and imaged offline (brightfield and fluorescence) using an IX70 Olympus inverted microscope (Olympus, Tokyo, Japan) with a 4x objective and a AmScope Microscope Digital Camera (MU1003, Amscope, Irvine, California, USA). Images of the MTs in each well were taken at T0 and after staining at T72.

### 2.14 Measurement of gaseous diffusion

Paramount to the success of mammalian cells’ viability is the ability of gases to diffuse through polymeric membranes. To ascertain gaseous diffusion through the adhesive polyurethane membrane and 1mm silicone sheet individually, an experiment was conducted by vacuum degassing the system for five minutes and measuring the amount of time taken to re-aerate the fluid. Utilising an optoelectrical oxygen sensor (PICO-O2, Pyro science, Aachen, Germany), phase-shift measurements were recorded to calculate the re-aeration rate of oxygen through the membranes.

### 2.15 Measurement of liquid diffusion

Blue food dye infused gelatin (4% w/v) was added to wells in the 3×3 prototype and left for 16hr at room temperature at 37°C. Images of diffusion were captured at 0 and 16h.

### 2.16 Biological Assays

#### 2.16.1 Albumin assay

Secretion of albumin was measured by Human Albumin Enzyme-Linked Immunosorbent Assay (ELISA) kit (Bethyl Laboratories, MA, USA). The ELISA assay was performed in clear flat-bottom 96-well microplates. Supernatant of HepG2 MTs cultured in 96 microplates were collected prior to feeding every 3-4 days and samples were stored frozen at -20°C prior to performing the assay. Prior to this, the assay samples were thawed to room temperature and prepared in accordance to the manufacturers protocol. Absorbance was measured at 450 nm at room temperature using a microplate reader (TECAN Infinite F nano, Männedorf, Switzerland).

A 4-parameter logistic (F (x) = d+ (a-d)/ (1+ (x/c)^b)) curve fit was performed on the ELISA measurement reading (MyAssays software). The level of albumin obtained from the supernatant of HepG2 microtissues were seen to increase from day 4 to 18.

#### 2.16.2 Urea assay

Urea assay was performed in accordance with the Urea assay kit (Sigma-Aldrich, Germany). In brief, the supernatant of HepG2 MTs cultured in 96 microplates were collected and stored at -20°C. Samples were diluted with the supplied buffer to adjust concentrations to the linear range of the assay. Assays were run in clear flat-bottom 96-well plates and measured at 570 nm using a microplate reader (TECAN Infinite F nano, Männedorf, Switzerland).

#### 2.16.3 CYP expression

HepG2 cells were seeded using a Multiflo FX multi-mode automated reagent dispenser (Biotek, UK) at 2.0 × 10^5^ cells per well (2D) in Ibidi microplate 96 well black plates (Ibidi, USA) and incubated for 24 h at 37 °C in a Forma Steri-Cycle CO2 incubator (Thermo Scientific, UK) or 400 cells per well (3D) in 96-well ultra-low attachment (ULA) round-bottomed plates (Corning, UK) in 180μL of complete DMEM. Cells were incubated for 4, 11 and 21 days 37 °C in a Forma Steri-Cycle CO_2_ incubator (Thermo Scientific, UK). Post-seeding CYP1A2 and CYP2E1 mRNA expression in HepG2 2D and 3D cultures were measured relying on RT-PCR. The results are expressed as percent mean mRNA content normalised to GAPDH and are the mean ± SD of n = 3 experiments.

#### 2.16.4 APAP toxicity in 3D HepG2 cultures in ULA μ-plate and Mera Bio Rig

HepG2 cells were seeded using a Multiflo FX multi-mode automated reagent dispenser (Biotek, UK) at 2.0 × 10^5^ cells per well (2D) in Ibidi 96 well black microplates (Ibidi, USA) and 200 cells per well (3D) in 96-well ultra-low attachment (ULA) round-bottomed plates (Corning, UK). Cells were incubated for 24 h (2D) and 7 days (3D) at 37 °C, 5% CO_2_. 2D and 3D cell cultures were treated with 30 mM of APAP for 24 h. Cells were subsequently stained with Calcein AM (1.5μM) and Ethidium homodimer-1 (10μM) and cell death and viability data were measured using Operetta CLS High Content Analysis System (PerkinElmer, UK).

For cells assessed on the Mera system, HepG2 cells were cultured in ULA microplates for 7 days and transferred to Mera for treatment and staining. 3D microtissues were stained with Calcein AM (3μM) and Ethidium homodimer-1 (10μM) on Mera and were then transferred to a ULA microplate for imaging consistency. Pixel intensity was measured using an IX70 Olympus inverted microscope (Olympus, Tokyo, Japan).

### 2.17 Statistical analysis

The acquired data is generally represented as mean values and their standard deviations.

## 3 Results and Discussion

The percentage of dissolved oxygen in the liquid media (% air saturation) was utilised to measure the gaseous diffusion through the polyurethane and silicone membranes. The phase shift measurements and oxygen levels follow an inverse relationship. The results in Figure 6 presents the re-aeration time for the polyurethane and silicone membrane of 1mm thickness was 9 minutes and 14 minutes, respectively. Based on the data generated, as the silicone was used to create a seal within the channel whilst maintaining the capability for gaseous diffusion, it was decided to use the silicone membrane instead of a combination of the silicone with the polyurethane membrane. This also reduces system complexity, increases accessibility, and reduces the amount of time required for experimental setup.

**Figure 6:**
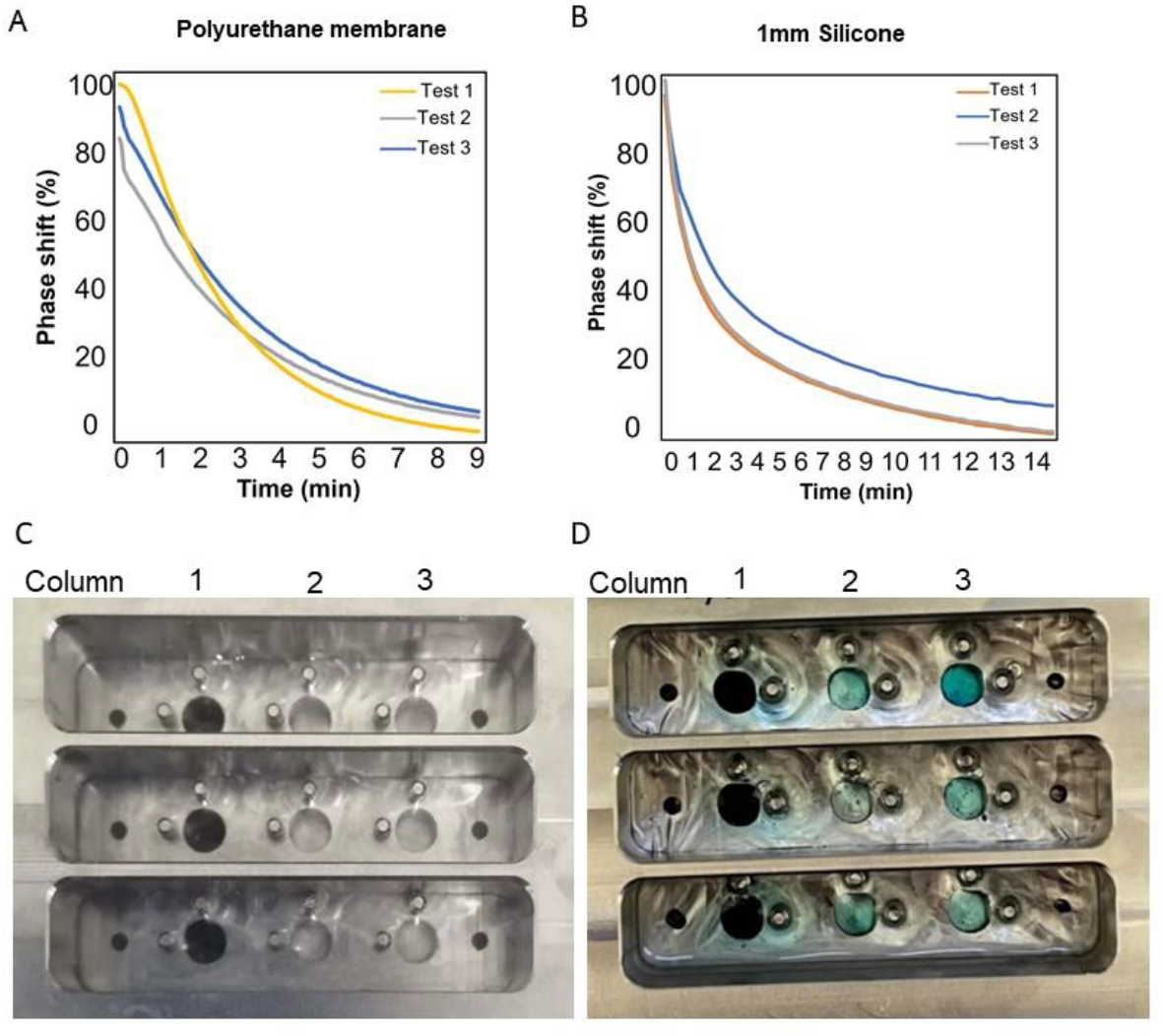
Measurement of diffusion in static condition, A) Oxygen diffusion through polyurethane membrane, B) Oxygen diffusion through a silicone membrane of 1mm thickness C) Demonstration of dye diffusion from wells in 3 × 3 Mera Bioplate at 0h and D) after 16h.

Figure 6 (C & D) presents a demonstration of liquid diffusion across well columns in the 3×3 prototype. Food dye infused 4% gelatin was added to wells in column 1 of the prototype and this set up was left for 16h at 37°C. The diffusion of dye out of the wells into the surrounding fluid was demonstrated over the time period as can be seen from the blue tint of the liquid around the wells in Figure 6, D. This shows the movement of small molecules (Brilliant Blue E133, 783 Da and Carmosine E122, 502 Da) out of the wells and into the surrounding fluid under static flow conditions. Variations in dye intensity in the wells can be noted which can be a result of small changes in the geometry of the channel increasing or decreasing diffusion rates. This demonstration is purely for visualisation purposes. A more rigid silicone layer can be employed in place of the more flexible polyurethane to offer more reproducible channel geometry. It can be theorised that under constant or pulsed perfusion recirculation of fluid may accelerate the diffusion process.

The objective of this paper was to demonstrate that the Mera system can culture and maintain/drug 3D microtissues for up to 72h and subsequently undertake staining to determine viability in an automated fashion. HepG2 was chosen as a liver model as it is an immortalised cell line that is low cost to purchase, easy to maintain, has high proliferation and retains many liver specific functions like albumin, urea and glucose secretions. However, it has limitations including low drug metabolizing phase I and phase II enzymes. Freshly isolated or cryopreserved primary human hepatocytes (pHHs) are regarded as gold standard for toxicity analysis as they reflect the properties and functions of liver cells *in vivo*. However fast de-differentiation occurs as soon as they are cultured *in vitro* in 2D while other drawbacks include limited supply from a single donor impacting reproducibility for repeated experiments and heterogeneity of samples between individuals making standardisation difficult (Kammerer, 2021).

Images of MTs were taken to observe potential morphological changes as shown in Figure 7, C where HepG2 MT size increases from 220μm in diameter on day 7 to over 1000μm in diameter by day 18. Albumin and urea expression are used as an indicator of metabolic activity of the cell and in healthy HepG2 microtissues, albumin and urea secretion rises as incubation time increases. It has been previously reported that higher levels of cytochrome P-450 activity were found via real-time polymerase chain reaction (RT-PCR) when hepatocytes were organized as 3D aggregates over 2D cell culture (Hsiao et al., 1999). Albumin and urea secretion was quantified from samples collected from the supernatant of microtissue-culture wells on Days 4, 7, 10, 14 and 18 and assayed according to the manufacturer’s protocol. This was to determine that the models of choice displayed high levels of liver-specific functions. As displayed in Figure 7A, a notable increase in albumin secretion is observed after 7 days of incubation. The data presented correlates with the values reported in the literature (Godoy et al., 2013, Hsiao et al., 1999). Urea secretion also increases after day 7 to approx. 340ng/mL (Figure 7B). Secretion of these metabolites are well defined physiological functions of hepatocytes *in vivo* and therefore the analysis of the production of these functional end points provides a measurement of liver-specific functionality in the current microtissue model (Godoy et al., 2013). These findings help to prove that culturing of cells in 3D provides a more representative model that summarises several *in-vivo* like characteristics that are not obvious in 2D monolayer cultures.

**Figure 7:**
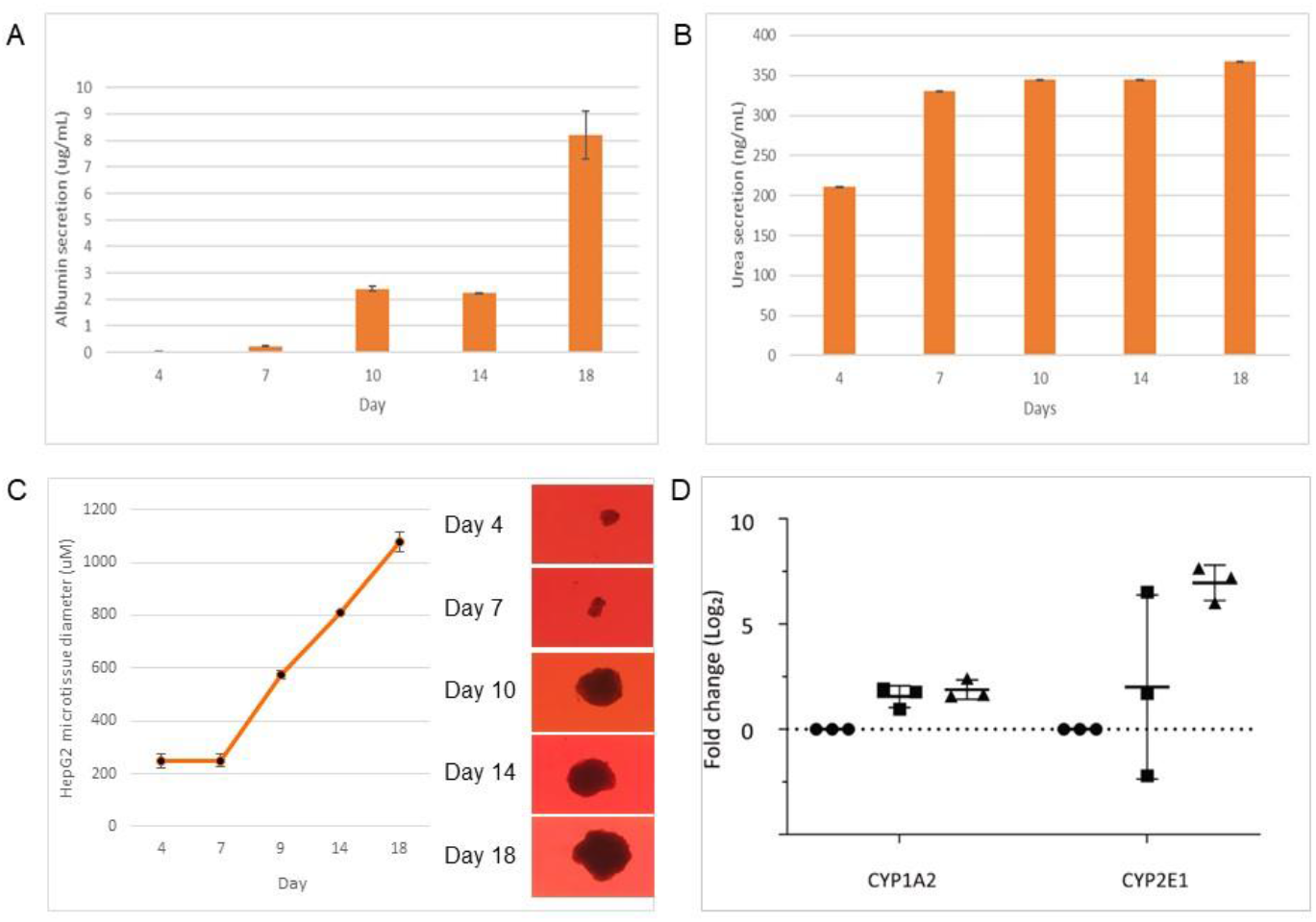
Characterisation of HepG2 microtissues (A) Albumin secretion concentrations (ug/mL) and (B) Urea secretion concentrations (ng/mL) from the supernatant of HepG2 microtissues from day 4-18. (n=3), (C) Growth profile of HepG2 microtissues over 18 days with sample images, n=3 (D) Expression of CYP1A2 and CYP2E1 in 2D and 3D HepG2 cell cultures. HepG2 cells were incubated for 4 days (2D) () or 11 (3D) () and 21 days () (3D) at 37 °C in a CO_2_ incubator. CYP1A2 and CYP2E1 mRNA expression in HepG2 2D and 3D cultures were measured using RT-PCR. The results are expressed as percent mean mRNA content normalised to GAPDH. (n = 3, *p* < 0.05).

Cytochrome P450 enzymes (CYPs) are involved in the oxidative metabolism of the majority of the commonly used low molecular weight drugs, thereby influencing the pharmacokinetics of these drugs and having an important role in drug-drug interactions. It is important that any cell models used to model drug interactions have sufficient levels of CYP expression to undertaken relevant drug metabolism. As displayed in Figure 7D, CYP expression in 3D cultures increased over 21 days in comparison to conventional 2D cell culture at day 4, particularly for CYP2E1 expression which increased over 5-fold over the time period. CYP levels in HepG2 cells in 2D culture are constant over time. Due to the growth rates of these cells in 2D culture the cells must be passaged every three to four days to avoid the cells becoming over confluent hence cell culture at 2D was limited at four days. 3D cultivation is seen as an important feature as it leads to increase in enzyme activity and hepatic function in comparison to 2D cell culture. When the HepG2 cells are induced to form 3D spheroids the local environment off the cells changes over the 21 days resulting in increased CYP expression. One of the recognised challenges with spheroid formation is reproducibility and this is especially evident at the 11 day intermediate time point where there is considerable variation in CYP expression. This variability stabilises by day 21.

APAP is a commonly used antipyretic and analgesic drug with a large therapeutic range, but at high doses or in combination with alcohol or other xenobiotics, it triggers centrilobular hepatic necrosis, resulting in acute liver failure. Approximately 5-10% of APAP is oxidised by CYP450s into N-acetyl-p-benzoquinone imine (NAPQI) which is detoxified with glutathione (GSH) upon conjugation. It is thought that the toxicity of APAP is directly linked to the binding of NAPQI to mitochondrial proteins (Lőrincz et al., 2021). To demonstrate the improved function of MTs, HepG2 cells grown in a 2D monolayer were compared to 3D MTs before and after treatment with 30mM APAP where APAP was used to induce cell death. Figure 8B display the results of this assessment with HepG2 MT viability significantly decreased after treatment at 2D. Images were taken with the Operetta CLS and image analysis undertaken using the Harmony software for area (um^2^) values for fluorescent images emitting at 520 nm for Calcein AM and 630 nm for Ethidium Bromide. The areas of the untreated samples were used to normalise the values for the treated samples by dividing the area of the treated samples by the area of the untreated samples. While this is a commonly used method for graphically representing this type of data especially where there are large changes in magnitude this method of normalisation can suppress variability. However, the levels of variability seen here are acceptable with the experimental parameters.

**Figure 8.**
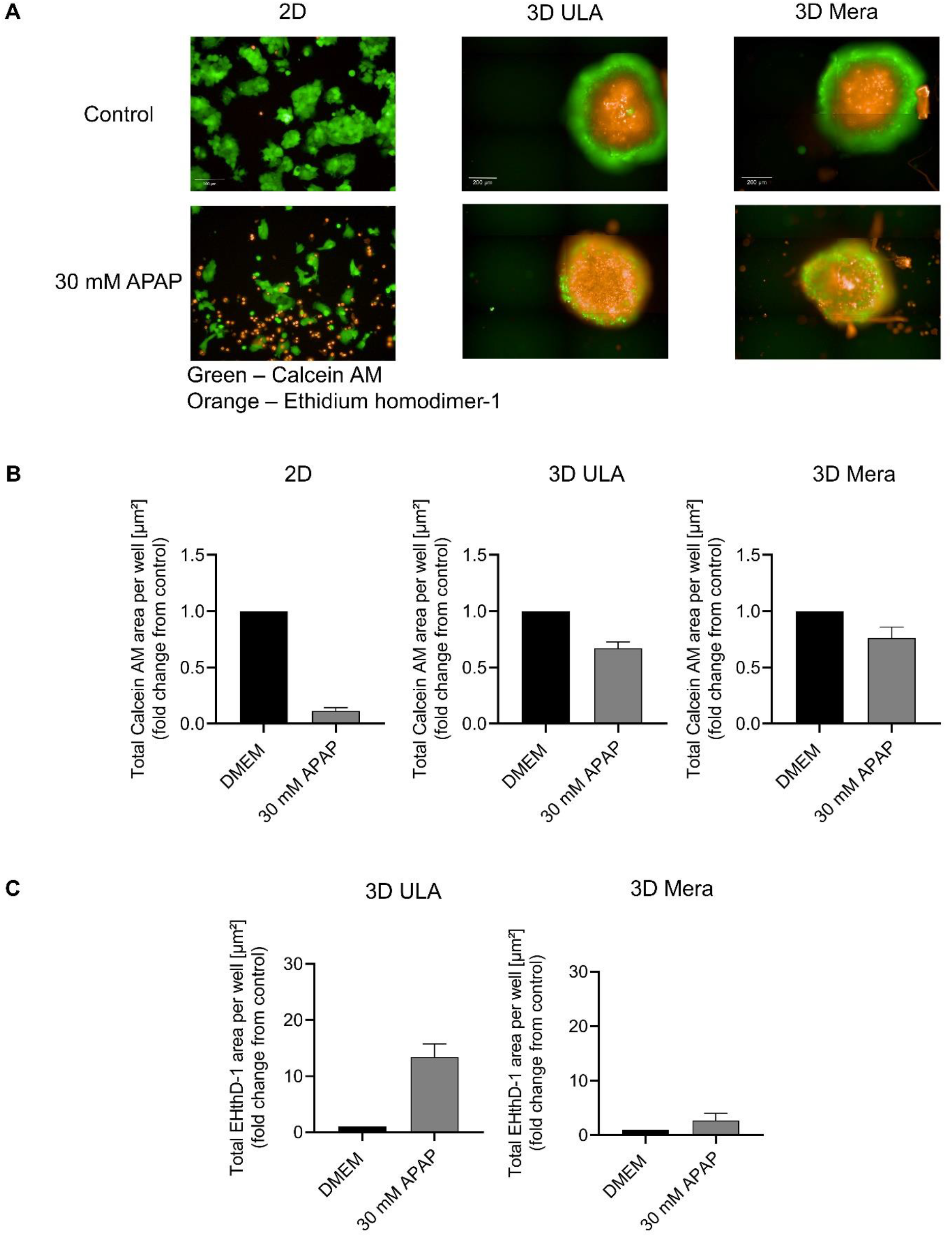
The effect of APAP toxicity in 3D HepG2 cultures in ULA microplate and Mera Bioplate (A) Representative images of 2D and 3D cell culture treatment groups. (B) Data are expressed as mean fold change in total area (μm^2^) of Calcein AM and (C) EtHD1 stained cells between treatment groups and controls (in 9 fields for 2D and the whole area of each spheroid) (± SD) of 3 independent experiments (1-5 replicates per treatment group). All images are the same magnification and shown at the same scale.

However reduced cell death is seen in 3D MT’s in ULA plates and in Mera after treatment inferring that their 3D structure provides a level of resistance against APAP toxicity. More interestingly, APAP is significantly less toxic to MTs in Mera than in the ULA plates which can be attributed to the frequent media change and perfusion flow in Mera. Also, the images of 3D in ULA microplates and 3D in Mera portray similar images suggesting that Mera provides an equivalent if not better environment to support MT growth and viability. From Figure 8, a necrotic core can be identified in all images. It is widely accepted that MT growth advances in 3 phases, phase 1 where cell cycling is observed throughout, phase 2 where MT’s develop to a size where cells in the core remain viable but enter cell cycle arrest and phase 3 where the MT develops a necrotic core (Browning et al., 2021). This is mainly due to limited oxygen and nutrient diffusion to the core as well as increases in carbon dioxide and waste products from the cells. This necrotic core is often seen in viable healthy MT’s and is not a cause for concern as long as the cells in the outer boundaries of the MT are alive and proliferating.

The Mera system allows us to perform both 3×3 and 4×10 Bioplate perfusion experiments to investigate the viability of HepG2 microtissues over several days. It was established that microtissues within the closed automated system which were stained online were viable for three days (Figure 9) with the expectation that this time period can be extended to 21 days and further. As described previously, serum free media was perfused for 1 minute every hour over the microtissues at a flow rate of 2.0 mL/min. The flow rate during medium circulation is adjustable. A concentration of 20mM APAP was selected to induce microtissue death over 72h. This concentration has previously been identified as optimal for induction of cell death by undertaking titrations of APAP diffusion up to 40mM on 96 well microplates on 2D and 3D cultures. Untreated microtissues significantly increased in size over the 72h period, from approx. 284uM ± 32.1 at 0h to 362uM ± 23.3 at 72h within the system. This suggests that uniform mixing of drugs occurs within the wells. This can be established to be true by the successful uniform staining of microtissues in Mera with Calcein AM which will not occur unless all media is removed from the wells and replaced with PBS. The images presented in Figure 9 are slightly blurred as imaging was undertaken on microtissues *in situ* within the machined acrylic well. Later prototypes will use injection moulded wells which will have improved clarity for imaging.

**Figure 9:**
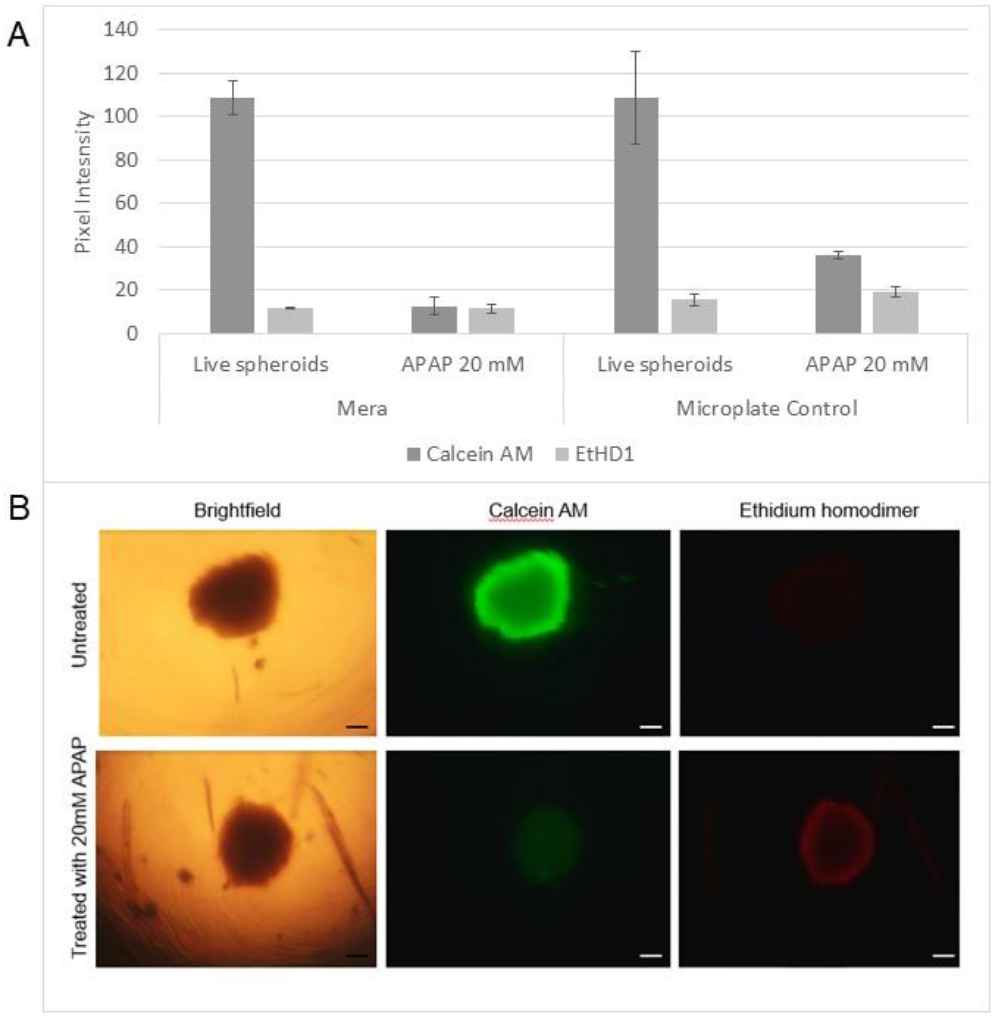
Evaluation of HepG2 microtissue viability in Mera (A) 3 × 3 Bioplate over 72 hours incubation. Microtissues cultured on Mera were stained with 5μM Calcein AM & 10μM EtHD1 while microtissues on the microplate were stained with 3μM Calcein AM & 10μM EtHD1, n=6. (B) Representative brightfield and fluorescence images of HepG2 microtissues treated with 20mM APAP and untreated in the Mera 3 × 3 Bioplate after 72 h incubation (scale bar = 100uM).

Online staining of HepG2 microtissues has successfully scaled on the 4×10 grid format with viability established after 24 hr culture as displayed in Figure 10, A. The ethidium homodimer stain has been optimised on the 4×10 Bioplate over 48h showing increased cell death after treatment with 20mM APAP. The Calcein AM viability stain is currently undergoing optimisation on the system and further data will be presented in subsequent publications.

**Figure 10:**
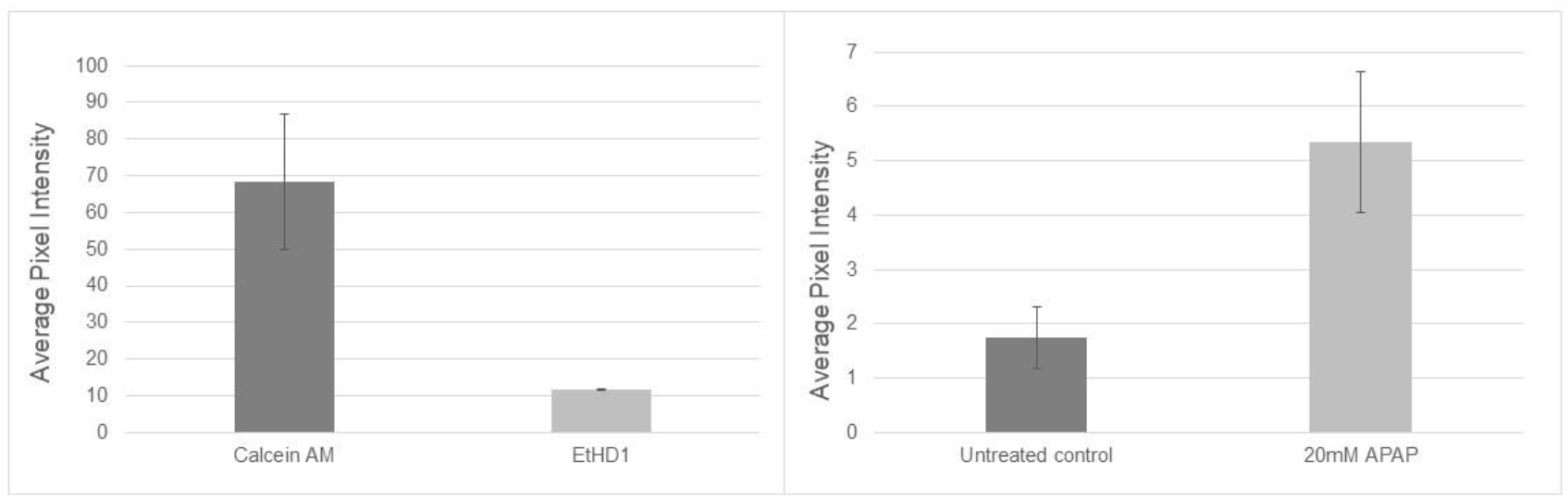
(A) Online staining on Mera 4 × 10 Bioplate after incubation of HepG2 microtissues over 24h. Viability was assessed by staining online in Mera with 5μM Calcein AM & 10μM EtHD1, n=39. (B) Evaluation of HepG2 microtissue control and treated with 20mM APAP on Mera 4×10 Bioplate over 48h incubation. Viability was assessed by staining online in Mera with 10μM EtHD1 alone, n=10.

As can be seen from the results, Mera consistently maintains healthy viable MTs for at least 72hrs in the 3×3 Bioplate format. The system demonstrates that the gas diffusion and liquid exchange are sufficient to support healthy growth of human microtissues. The APAP based toxicity assay also demonstrates the system’s capacity for modelling drug interaction and online viability testing. These parameters have been demonstrated in the 3×3 prototype and in the 4×10 prototype. It is envisaged that the latter will be the basic unit size of the modular Mera system with scaling being facilitated through replicates of the 4×10 module. The system will also be tested with lengthier runs in the timeframe of weeks, with the aim of marketing a MPS that can run automated 3D assays for up to 21 days.

Mera has several advantages over current MPS on the market. The system was designed to operate without the use of PDMS which is frequently used in MPS and Organ-on-a-chip designs despite the fact it has high absorption for small hydrophobic molecules. All materials used in Mera were selected not only for their stability and durability but also for their biocompatibility and compatibility with commercial sterilisation protocols as determined from thorough assessment of each materials technical specifications. A key factor in the development in MPS’s is the need for systems that operate at the scale the drug developers require, often being able to assess 10’s or 100’s of drug candidates in parallel. When scaled, one assembled Mera platform is envisaged to be able to culture up to 800 microtissues concurrently whereby 20 modular SBS compatible 4×10 grid of wells (Bioplates) will be accommodated in one comprehensive system. The proposed commercial system design includes 20 modules, as shown in Figure 12 within a housing unit of approx. 1m^3^ dimensions, which would allow Mera to be marketed a desktop device. Where desired the system could be scaled beyond this size. This is up to 10X more throughput than currently available microphysiological systems or perfused OoC systems on the market. For example, CN-Bio offers the PhysioMimix organ-on-a-chip perfusion system provided in single-organ and multi-organ formats and specifically provide a Liver plate for 3D coculture of primary human liver cells (which also can accommodate induced pluripotent stem cells, immortalised cells lines and tissue slices) of up to 4 weeks duration. However, the PhysioMimix system has limited throughput with plates containing up to 12 wells per plate and only up to 6 plates can be run in parallel (CN Bio Innovations, 2022).

Although a silicone gasket was used in conjunction with a polyurethane membrane to maintain sealing integrity and capability for gaseous diffusion for the data generated presented in this paper, it was decided to eliminate use of the polyurethane membrane for future experiments. The gaseous diffusibility of oxygen though 1mm silicone provide enough oxygen to support the high oxygen demands of HepG2 microtissues (Sharifi et al., 2019). Removal of the polyurethane sheet also reduces system complexity, increases accessibility, and reduces the amount of time required for experimental setup. The Bioplate will be redesigned to ensure compatibility with ANSI/SBS 4-2004 standards for formats used with plate handling technology. In the future, the Bioplate will be manufactured via injection moulding using a polymeric material such as polystyrene (PS) to decrease production time and increase the optical clarity when imaging (Campbell et al., 2021). Bioplates supplied with the system will be single use and provided in a sterile state. The next iteration of the system is being designed to lower the amount of fluid required for cost efficiency and to more closely mimic the physiological ratio of drugs and analytes to fluid (Malik et al., 2021). The Bioplates can also be removed from Mera and used in conjunction with standard microscopy equipment if required.

As discussed previously, dye diffusion experiments have demonstrated that under static conditions, diffusion will occur within the system while the viability of the microtissues over 72 hours proves nutrient diffusion has occurred. Up to four different organ types can be accommodated on the 4×10 Bioplate as per user requirement (e.g. gastrointestinal, liver, kidney and heart as shown in Figure 1,C) with each row of ten housing a different microtissues type. Flow can be directed across rows of a common tissue type (Y-direction) for growth or maintenance or alternatively flow can be switched to direct flow along a channel of four different tissue types (X-direction) thereby initiating recirculation flow to facilitate interorgan communication. Preliminary data to assess the effect of recirculation on the viability of HepG2 over 48 hours was generated as shown in Figure 11. There is a decrease in the pixel intensity of the MTs grown under static conditions in Mera in comparison to those that were exposed to perfusion over 48h (Figure 11, A). This can also be observed from the images in Figure 11, B where an obvious decrease in Calcein AM signal and an increase in EtHD1 can be seen. These are promising observations particularly for the use of Mera for intertissue communication and future publications will report further investigations generated from the recirculation system (described in Fig. 3) in greater detail. Peristaltic pumps displace fluid by mechanically actuating a flexible channel structure. There are a number of advantages to the use of peristaltic pumps over more commonly used gravity driven systems including tunable flow rates, unidirectional fluid flow and the ability to work with low fluid volumes (Schneider et al., 2021). Studies have been undertaken to assess pump-free, tilt based medium recirculation however such systems lack precise flow modification and large fluid volumes are required (Lohasz et al., 2018, Sasserath et al., 2020). The introduction of the closed-loop recirculation increases the opportunity for intercellular communication through frequent perfusion across the different cell types of the same fluid batch. Ratios of nutrient replacement, tissue mass to fluid volumes will be considered and designed to not affect the feeding regime and avoid removal of any molecules essential to inter tissue communication. This is currently in the validation stage. The design of the system was carefully evaluated to maintain sterility of the system. Mera is a sealed system, thereby not only avoiding evaporative fluid loss but also reducing contamination points. The current disinfection cycle is operating successfully, using 70% IPA and PBS and takes approximately twenty minutes per cycle to clean each line individually. The Bioplate will be provided in a sterile state and will be single use requiring replacement after each experiment to ensure sterility of the whole system. Material selection is key to this process for example, polyetheretherketone (PEEK) will replace many key components of the valving system as it is autoclavable and its use demonstrated in many biomedical designs (Kumar et al., 2018).

**Figure 11:**
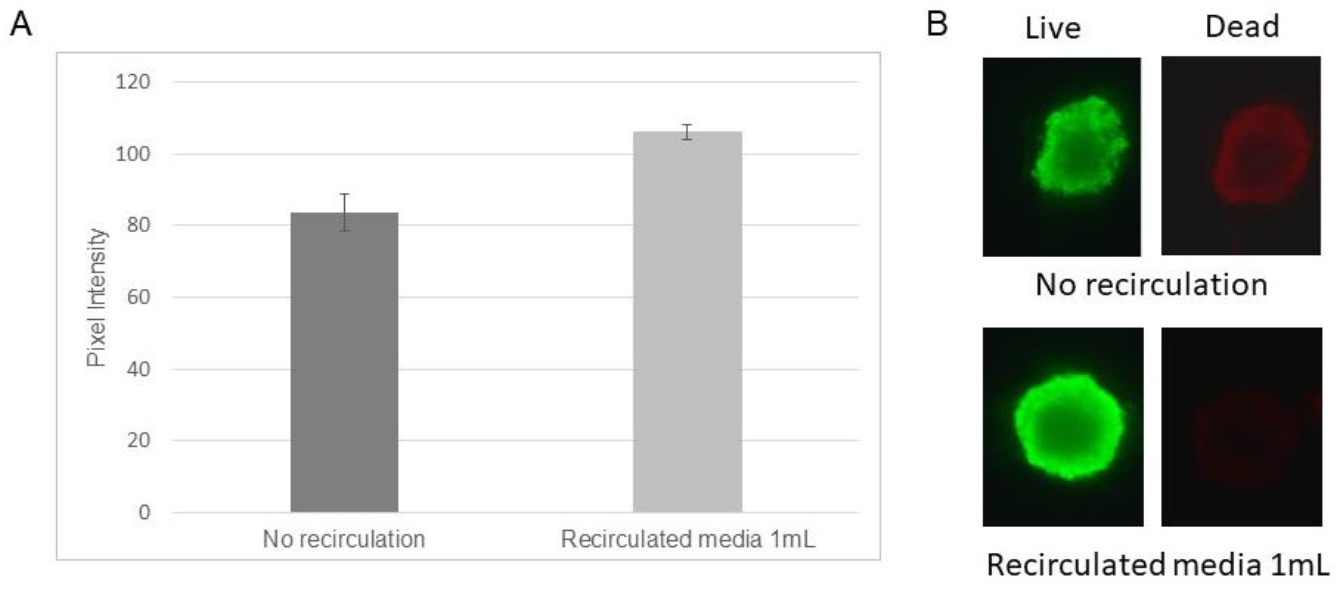
A) Assessment of HepG2 microtissues in static culture in Mera 3 × 3 format (no recirculation) versus recirculated media (1mL/hour) over a 48-hour period, B) Comparison of microtissues in Mera under static versus recirculated media over 48h. Microtissues were stained with 5μM Calcein AM & 10μM EtHD1, n=3.

**Figure 12:**
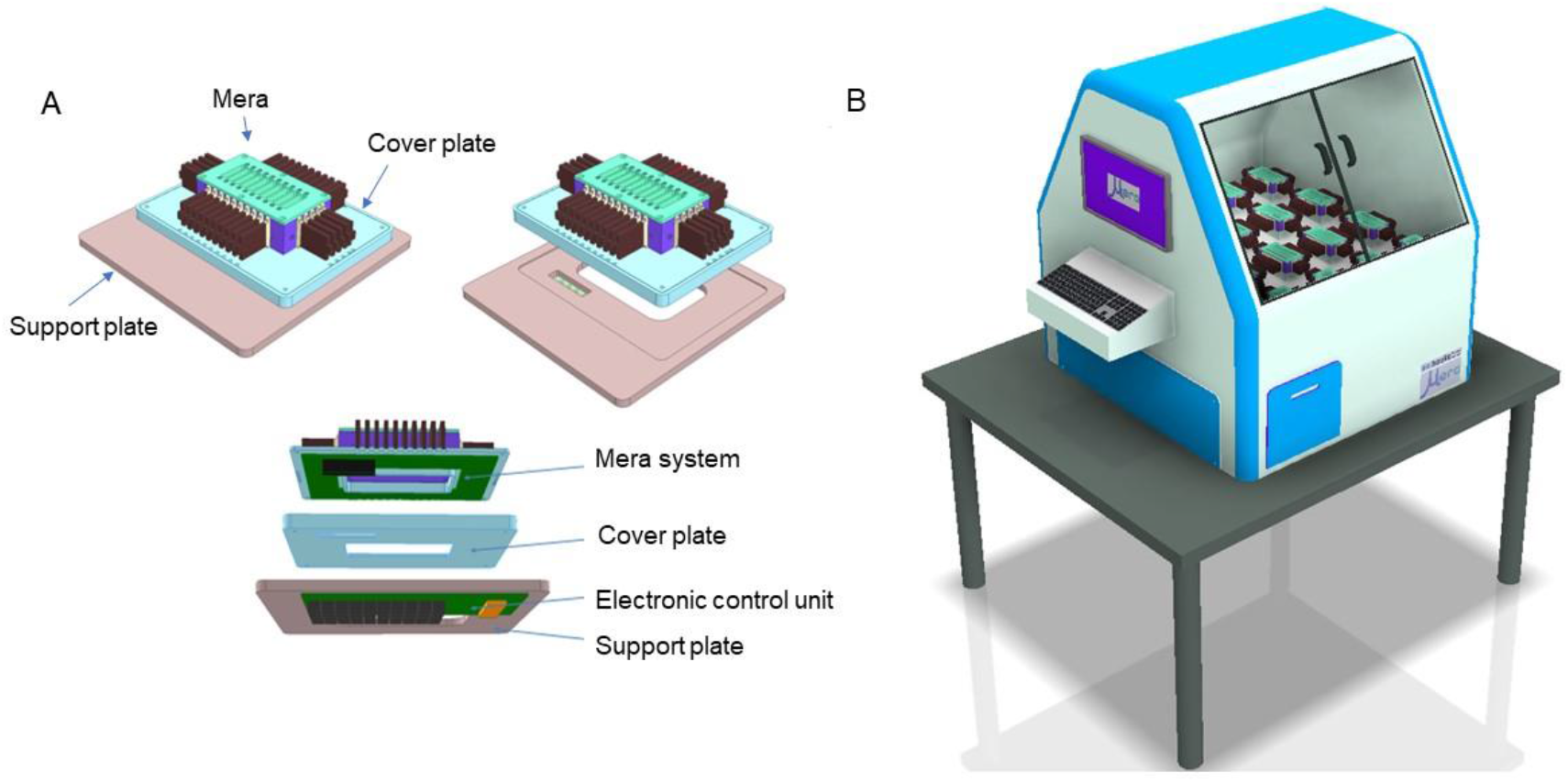
A) Proposed assembly of the 4 × 10 Bioplate and electronic control module, B) Proposed commercial Mera system (1m^3^ dimensions) showing individual fluidic modules (20 modules) within the unit.

Socio-political pressure requires the pharma industry to embrace more humane and ethical R&D approaches using cost-effective and reliable methodologies. European initiatives such as the 3R’s and the Animal Research: Reporting of *in vivo* Experiments (ARRIVE) guidelines (NC3Rs, 2022) have been adopted by the European Medicines Agency (EMA). While in the USA, the FDA, Defence Advanced Research Projects Agency (DARPA), National Institutes of Health (NIH) MPS programme (Sutherland et al., 2013) and the MPS consortium Cooperative Research and Development Agreement (CRADA) have employed similar initiatives. Groups composed of well-respected industry and academic advisors such as Human Organ and Disease Model Technologies (hDMT)(hDMT, 2022), Organ-on-a-chip in Development (ORCHID)(European Commission, 2022) and European Organ-on-chip (EUROoC)(EUROoC, 2022) are also involved in promoting regulatory guidelines and standardisation for MPS in the EU. High throughput based MPS with the ability to culture multi-organ systems offer promise for reducing the need for animal studies particularly for use in preclinical and early-stage clinical trials. The reduction and perhaps eventual replacement of preclinical animal models with MPS technologies and their integration into drug development pipelines and inclusion of data generated using such models in FDA regulatory submissions is a complex issue. To reach this goal, pharmaceutical companies, and regulatory agencies will require robust evidence of equivalent or superior performance relative to animal models. In addition, the ability to culture patient derived MTs in Mera would enable utilisation of this system for personalised and precision medicine applications.

One of the main advantages of using Mera as a system is the ability to see how a drug and its subsequent reactants and analytes will affect other cells in the body, saving time and money on preclinical studies and reducing the need for animal testing. A recent evaluation of MPS technologies concluded that if implemented into pharmaceutical R&D, up to 26% of R&D costs can be reduced as well as the turnover time in preclinical phases (Franzen et al., 2019). Its ability to tune perfusion and image in an automated fashion along with its modular design is beneficial for scaling and 3D cell culture. Long term culture of microtissues is desirable to provide the opportunity to analyse multiorgan interactions in a biomimetic microenvironment. Future work with Mera aims to extend the culture of HepG2 microtissues to up to 21 days at higher throughput than existing technologies on the market. We aim to investigate the functionality of MT’s on the Bioplate in respect to albumin, urea and CYP expression. We also aim to integrate patient derived samples or hiPSC in future experimental work which have higher CYP expression and albumin release. In addition, connection of multiple organ types via perfusion channels which can be changed as per the users’ requirements will be established to demonstrate interorgan communication and viability.

## Acknowledgements

The authors acknowledge financial assistance of Enterprise Ireland, the Western Development Commission, Boole Investment Syndicate, Irrus Investments, and the Disruptive Technologies Innovation Fund (Project no.: 164610/RR) who funded this research. The technology described in this article is in development and represents ongoing research and development efforts.

## Supplementary information

**Supplementary Figure 1:**
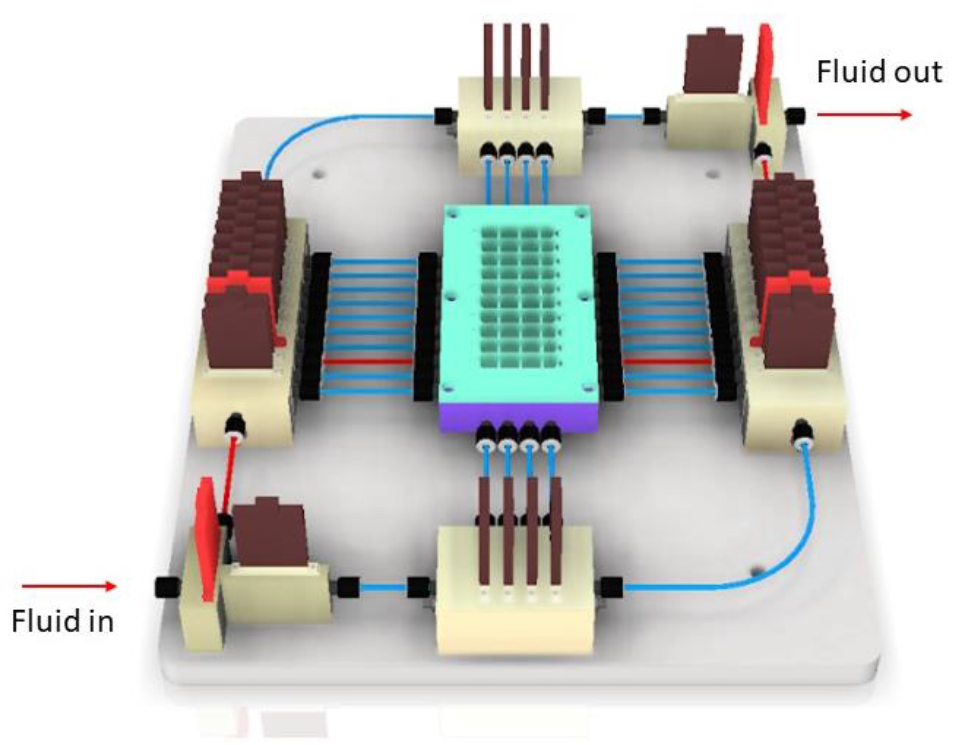
A) Fluid flow direction example (in red) through Mera system

**Supplementary Figure 2:**
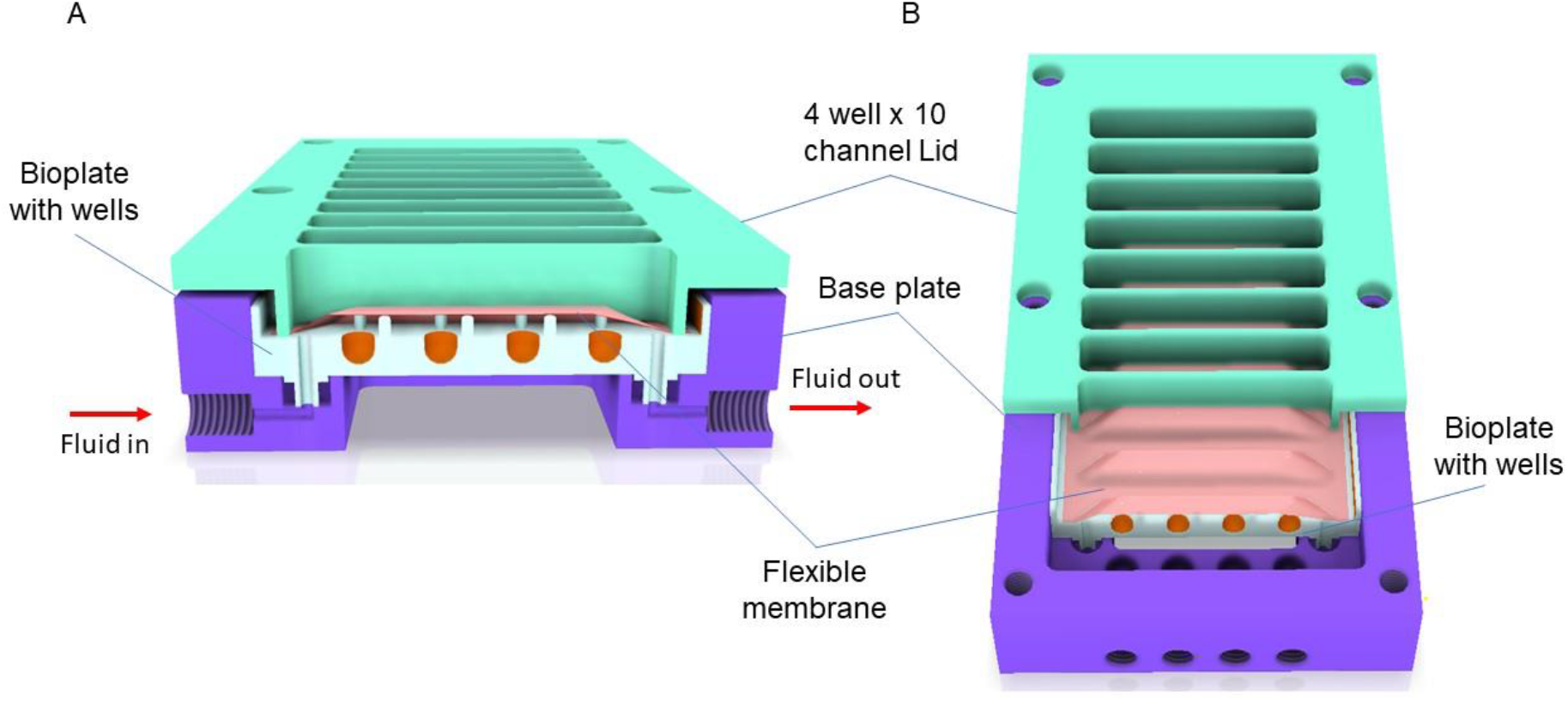
A) View X cutaway of 4 well x 10 channel lid and Bioplate assembly with flexible membrane, B) Top view of cutaway 4 well x 10 channel lid and Bioplate assembly with flexible membrane.

**Supplementary Figure 3:**
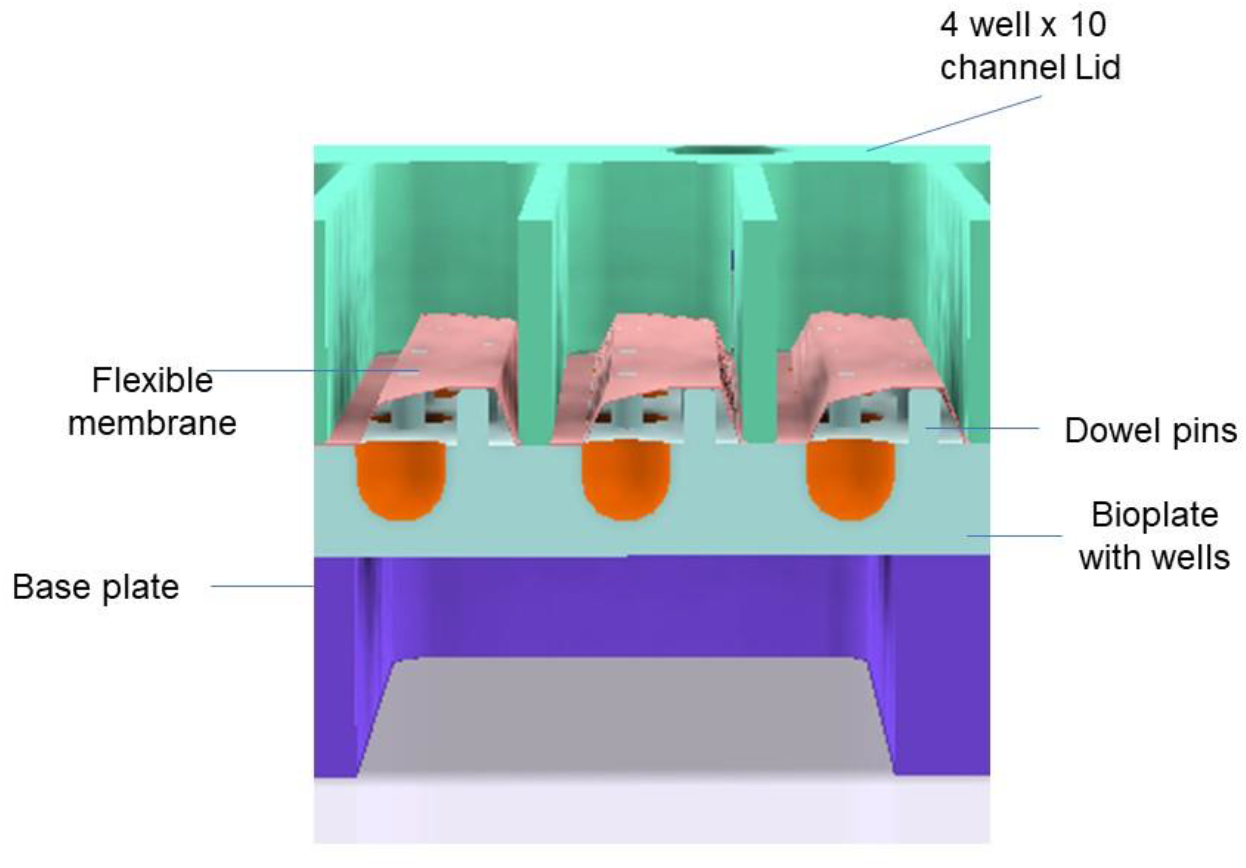
A) Alternative view Y cutaway of 4 well x 10 channel lid and Bioplate assembly with flexible membrane

**Supplementary Figure 4:**
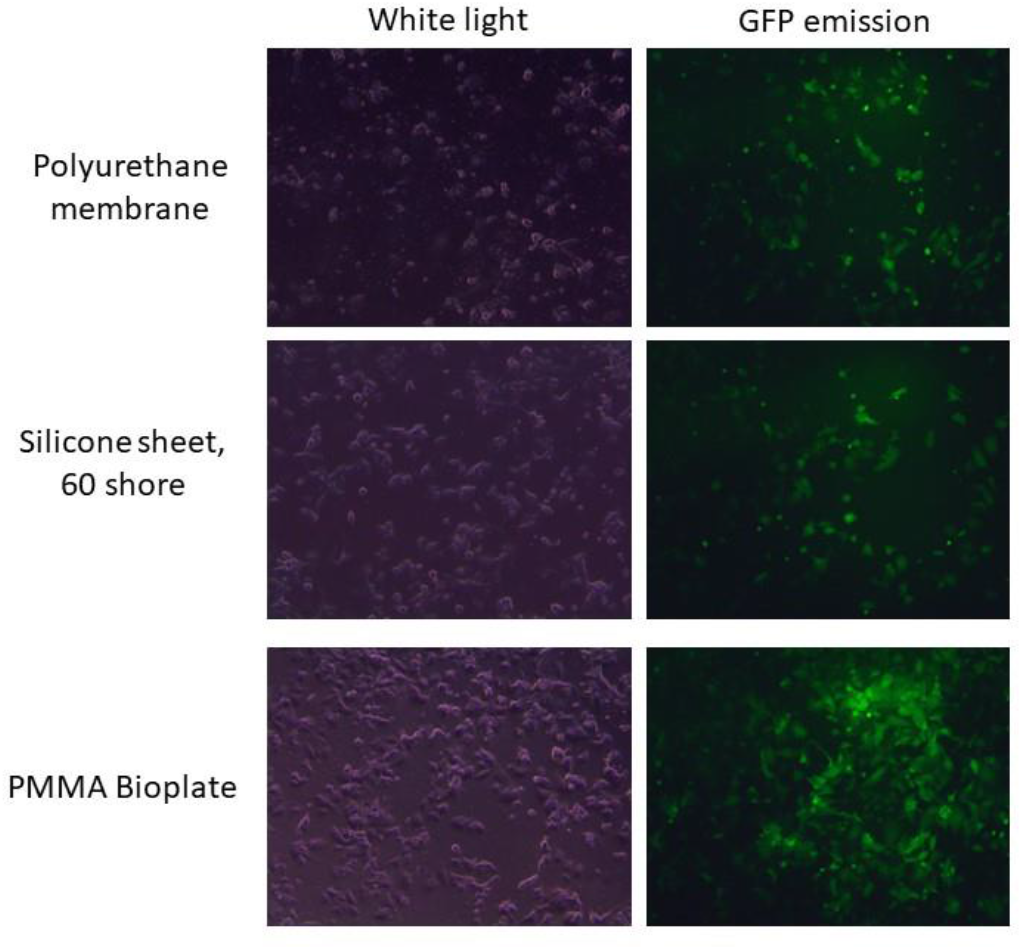
Images of using MCF7-GFP cell line exposed to Mera component materials after 7 days. Cells were plated in 2D on a 12 well microplate containing a 1cm X 1cm sample of the material of interest and imaged using Olympus IX70 fluorescent microscope.

**Supplementary Figure 5:**
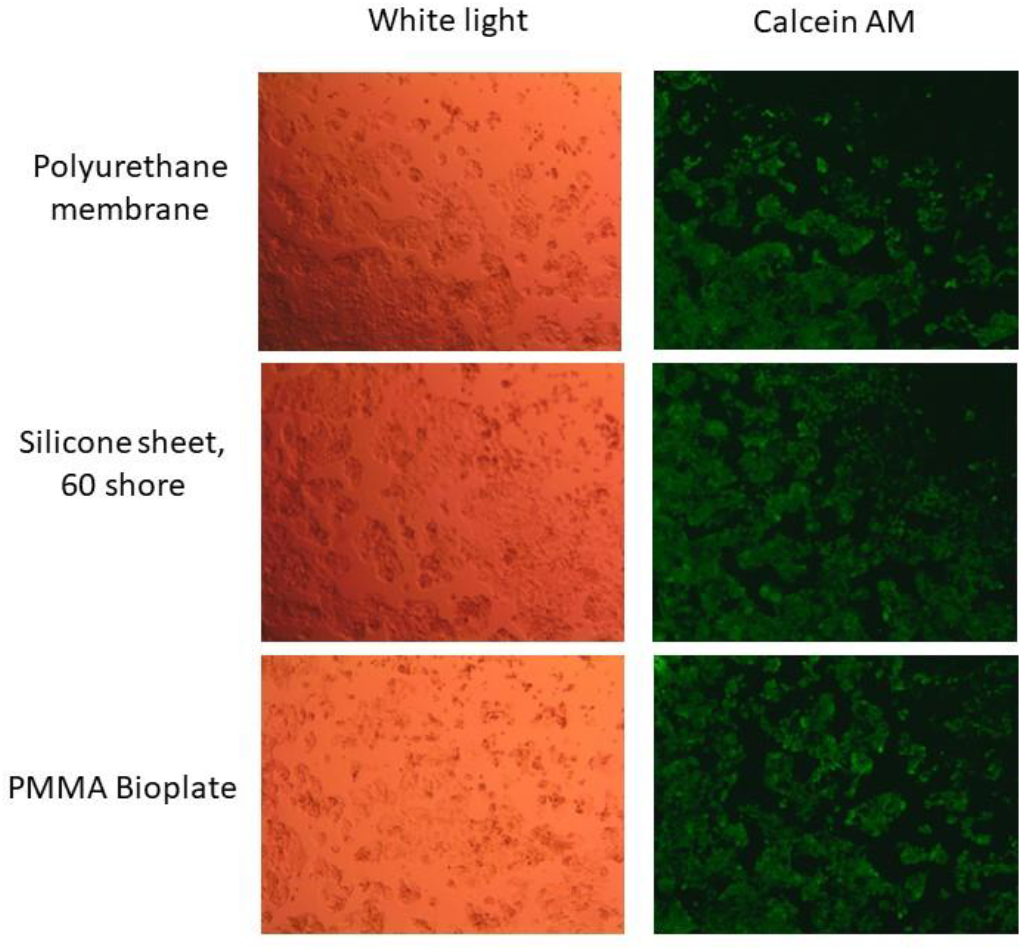
Images of using HepG2 cell line exposed to Mera component materials after 7 days. Cells were plated in 2D on a 12 well microplate plate containing a 1cm X 1cm sample of the material of interest, stained using 1.5uM Calcein AM and imaged using an Olympus IX70 fluorescent microscope.

All materials in contact with cells within the system were tested for biocompatibility by incubating HepG2 and MCF-GFP cells for up to 7 days in the presence of the material of interest and cell morphology and viability was subsequently assessed. All material samples were sterilised prior to placing into the 12 well microplate by treating firstly with 70% IPA and secondly with UV irradiation for 15 min to ensure sterility.

## Footnotes

1 Microphysiological systems (MPSs), Microtissue (MT), Polyetheretherketone (PEEK), Polytetrafluoroethylene (PTFE), Polystyrene (PS)

## References

Ahadian, S., Civitarese, R., Bannerman, D., Mohammadi, M. H., Lu, R., Wang, E., Davenport-Huyer, L., Lai, B., Zhang, B., Zhao, Y., Mandla, S., Korolj, A. & Radisic, M. 2018. Organ-On-A-Chip Platforms: A Convergence of Advanced Materials, Cells, and Microscale Technologies. Advanced Healthcare Materials, 7.

Ainslie, G. R., Davis, M., Ewart, L., Lieberman, L. A., Rowlands, D. J., Thorley, A. J., Yoder, G. & Ryan, A. M. 2019. Microphysiological lung models to evaluate the safety of new pharmaceutical modalities: a biopharmaceutical perspective. Lab Chip, 19, 3152–3161.

Asif, A., Park, S. H., Manzoor Soomro, A., Khalid, M. A. U., Salih, A. R. C., Kang, B., Ahmed, F., Kim, K. H. & Choi, K. H. 2021. Microphysiological system with continuous analysis of albumin for hepatotoxicity modeling and drug screening. Journal of Industrial and Engineering Chemistry, 98, 318–326.

Berger, D. R., Ware, B. R., Davidson, M. D., Allsup, S. R. & Khetani, S. R. 2015. Enhancing the functional maturity of induced pluripotent stem cell-derived human hepatocytes by controlled presentation of cell-cell interactions in vitro. Hepatology, 61, 1370–81.

Browning, A. P., Sharp, J. A., Murphy, R. J., Gunasingh, G., Lawson, B., Burrage, K., Haass, N. K. & Simpson, M. 2021. Quantitative analysis of tumour spheroid structure. eLife, 10, e73020.

Campbell, S. B., Wu, Q., Yazbeck, J., Liu, C., Okhovatian, S. & Radisic, M. 2021. Beyond Polydimethylsiloxane: Alternative Materials for Fabrication of Organ-on-a-Chip Devices and Microphysiological Systems. ACS Biomaterials Science & Engineering, 7, 2880–2899.

Chen, P.-Y., Hsieh, M.-J., Liao, Y.-H., Lin, Y.-C. & Hou, Y.-T. 2021. Liver-on-a-chip platform to study anticancer effect of statin and its metabolites. Biochemical Engineering Journal, 165, 107831.

Cn Bio Innovations. 2022. CN-BIO [Online]. Available: https://cn-bio.com/ [Accessed 11/10/2022 2022].

Conant, G., Lai, B. F. L., Lu, R. X. Z., Korolj, A., Wang, E. Y. & Radisic, M. 2017. High-Content Assessment of Cardiac Function Using Heart-on-a-Chip Devices as Drug Screening Model. Stem Cell Rev Rep, 13, 335–346.

Ding, E. L., Song, Y., Manson, J. E., Hunter, D. J., Lee, C. C., Rifai, N., Buring, J. E., Gaziano, J. M. & Liu, S. 2009. Sex Hormone–Binding Globulin and Risk of Type 2 Diabetes in Women and Men. New England Journal of Medicine, 361, 1152–1163.

Edington, C. D., Chen, W. L. K., Geishecker, E., Kassis, T., Soenksen, L. R., Bhushan, B. M., Freake, D., Kirschner, J., Maass, C., Tsamandouras, N., Valdez, J., Cook, C. D., Parent, T., Snyder, S., Yu, J., Suter, E., Shockley, M., Velazquez, J., Velazquez, J. J., Stockdale, L., Papps, J. P., Lee, I., Vann, N., Gamboa, M., Labarge, M. E., Zhong, Z., Wang, X., Boyer, L. A., Lauffenburger, D. A., Carrier, R. L., Communal, C., Tannenbaum, S. R., Stokes, C. L., Hughes, D. J., Rohatgi, G., Trumper, D. L., Cirit, M. & Griffith, L. G. 2018. Interconnected Microphysiological Systems for Quantitative Biology and Pharmacology Studies. Scientific Reports, 8, 4530.

Emulate Bio. 2022. Available: https://emulatebio.com/ [Accessed 2022].

EUROOC. 2022. EUROoC Network [Online]. Available: https://euroocs.eu/ [Accessed 08 August 2022].

European Commission. 2022. Organ on Chip in Development [Online]. Available: https://h2020-orchid.eu/ [Accessed 16 August 2022].

Ewart, L., Apostolou, A., Briggs, S. A., Carman, C. V., Chaff, J. T., Heng, A. R., Jadalannagari, S., Janardhanan, J., Jang, K.-J., Joshipura, S. R., Kadam, M. M., Kanellias, M., Kujala, V. J., Kulkarni, G., Le, C. Y., Lucchesi, C., Manatakis, D. V., Maniar, K. K., Quinn, M. E., Ravan, J. S., Rizos, A. C., Sauld, J. F. K., Sliz, J. D., Tien-Street, W., Trinidad, D. R., Velez, J., Wendell, M., Irrechukwu, O., Mahalingaiah, P. K., Ingber, D. E., Scannell, J. W. & Levner, D. 2022. Qualifying a human Liver-Chip for predictive toxicology: Performance assessment and economic implications. bioRxiv, 2021.12.14.472674.

Franzen, N., Van Harten, W. H., RetÈl, V. P., Loskill, P., Van Den Eijnden-Van Raaij, J. & m, I. J. 2019. Impact of organ-on-a-chip technology on pharmaceutical R&D costs. Drug Discov Today, 24, 1720–1724.

Godoy, P., Hewitt, N. J., Albrecht, U., Andersen, M. E., Ansari, N., Bhattacharya, S., Bode, J. G., Bolleyn, J., Borner, C., BÖTtger, J., Braeuning, A., Budinsky, R. A., Burkhardt, B., Cameron, N. R., Camussi, G., Cho, C. S., Choi, Y. J., Craig Rowlands, J., Dahmen, U., Damm, G., Dirsch, O., Donato, M. T., Dong, J., Dooley, S., Drasdo, D., Eakins, R., Ferreira, K. S., Fonsato, V., Fraczek, J., Gebhardt, R., Gibson, A., Glanemann, M., Goldring, C. E., GÓMez-LechÓN, M. J., Groothuis, G. M., Gustavsson, L., Guyot, C., Hallifax, D., Hammad, S., Hayward, A., HÄUssinger, D., Hellerbrand, C., Hewitt, P., Hoehme, S., HolzhÜTter, H. G., Houston, J. B., Hrach, J., Ito, K., Jaeschke, H., Keitel, V., Kelm, J. M., Kevin Park, B., Kordes, C., Kullak-Ublick, G. A., Lecluyse, E. L., Lu, P., Luebke-Wheeler, J., Lutz, A., Maltman, D. J., Matz-Soja, M., Mcmullen, P., Merfort, I., Messner, S., Meyer, C., Mwinyi, J., Naisbitt, D. J., Nussler, A. K., Olinga, P., Pampaloni, F., Pi, J., Pluta, L., Przyborski, S. A., Ramachandran, A., Rogiers, V., Rowe, C., Schelcher, C., Schmich, K., Schwarz, M., Singh, B., Stelzer, E. H., Stieger, B., StÖBer, R., Sugiyama, Y., Tetta, C., Thasler, W. E., Vanhaecke, T., Vinken, M., Weiss, T. S., Widera, A., Woods, C. G., Xu, J. J., Yarborough, K. M. & Hengstler, J. G. 2013. Recent advances in 2D and 3D in vitro systems using primary hepatocytes, alternative hepatocyte sources and non-parenchymal liver cells and their use in investigating mechanisms of hepatotoxicity, cell signaling and ADME. Arch Toxicol, 87, 1315–530.

GÖRgens, C., Ramme, A. P., Guddat, S., Schrader, Y., Winter, A., Dehne, E.-M., Horland, R. & Thevis, M. 2021. Organ-on-a-chip: Determine feasibility of a human liver microphysiological model to assess long-term steroid metabolites in sports drug testing. Drug Testing and Analysis, 13, 1921–1928.

Hdmt. 2022. Human Organ and Disease Model Technologies [Online]. Available: https://www.hdmt.technology/ [Accessed 10 August 2022].

Herland, A., Maoz, B. M., Das, D., Somayaji, M. R., Prantil-Baun, R., Novak, R., Cronce, M., Huffstater, T., Jeanty, S. S. F., Ingram, M., Chalkiadaki, A., Benson Chou, D., Marquez, S., Delahanty, A., Jalili-Firoozinezhad, S., Milton, Y., Sontheimer-Phelps, A., Swenor, B., Levy, O., Parker, K. K., Przekwas, A. & Ingber, D. E. 2020. Quantitative prediction of human pharmacokinetic responses to drugs via fluidically coupled vascularized organ chips. Nature Biomedical Engineering, 4, 421–436.

Hofer, M. & Lutolf, M. P. 2021. Engineering organoids. Nature Reviews Materials, 6, 402–420.

Hsiao, C. C., Wu, J. R., Wu, F. J., Ko, W. J., Remmel, R. P. & Hu, W. S. 1999. Receding cytochrome P450 activity in disassembling hepatocyte spheroids. Tissue Eng, 5, 207–21.

Jalili-Firoozinezhad, S., Gazzaniga, F. S., Calamari, E. L., Camacho, D. M., Fadel, C. W., Bein, A., Swenor, B., Nestor, B., Cronce, M. J., Tovaglieri, A., Levy, O., Gregory, K. E., Breault, D. T., Cabral, J. M. S., Kasper, D. L., Novak, R. & Ingber, D. E. 2019. A complex human gut microbiome cultured in an anaerobic intestine-on-a-chip. Nature Biomedical Engineering, 3, 520–531.

Kammerer, S. 2021. Three-Dimensional Liver Culture Systems to Maintain Primary Hepatic Properties for Toxicological Analysis In Vitro. Int J Mol Sci, 22.

Kim, J., Koo, B.-K. & Knoblich, J. A. 2020. Human organoids: model systems for human biology and medicine. Nature Reviews Molecular Cell Biology, 21, 571–584.

Kumar, A., Yap, W. T., Foo, S. L. & Lee, T. K. 2018. Effects of Sterilization Cycles on PEEK for Medical Device Application. Bioengineering (Basel), 5.

Lancaster, M. A., Renner, M., Martin, C.-A., Wenzel, D., Bicknell, L. S., Hurles, M. E., Homfray, T., Penninger, J. M., Jackson, A. P. & Knoblich, J. A. 2013. Cerebral organoids model human brain development and microcephaly. Nature, 501, 373–379.

Li, C. Y., Cao, C. Z., Xu, W. X., Cao, M. M., Yang, F., Dong, L., Yu, M., Zhan, Y. Q., Gao, Y. B., Li, W., Wang, Z. D., Ge, C. H., Wang, Q. M., Peng, R. Y. & Yang, X. M. 2010. Recombinant human hepassocin stimulates proliferation of hepatocytes in vivo and improves survival in rats with fulminant hepatic failure. Gut, 59, 817–26.

Lind, J. U., Yadid, M., Perkins, I., O’Connor, B. B., Eweje, F., Chantre, C. O., Hemphill, M. A., Yuan, H., Campbell, P. H., Vlassak, J. J. & Parker, K. K. 2017. Cardiac microphysiological devices with flexible thin-film sensors for higher-throughput drug screening. Lab on a Chip, 17, 3692–3703.

Liu, L., Koo, Y., Akwitti, C., Russell, T., Gay, E., Laskowitz, D. T. & Yun, Y. 2019. Three-dimensional (3D) brain microphysiological system for organophosphates and neurochemical agent toxicity screening. PLoS One, 14, e0224657.

Lohasz, C., Rousset, N., Renggli, K., Hierlemann, A. & Frey, O. 2018. Scalable Microfluidic Platform for Flexible Configuration of and Experiments with Microtissue Multiorgan Models. SLAS TECHNOLOGY: Translating Life Sciences Innovation, 24, 79–95.

LŐRincz, T., DeÁK, V., Makk-Merczel, K., Varga, D., HajdinÁK, P. & Szarka, A. 2021. The Performance of HepG2 and HepaRG Systems through the Glass of Acetaminophen-Induced Toxicity. Life (Basel), 11.

Ma, C., Peng, Y., Li, H. & Chen, W. 2021. Organ-on-a-Chip: A New Paradigm for Drug Development. Trends Pharmacol Sci, 42, 119–133.

Malik, M., Yang, Y., Fathi, P., Mahler, G. J. & Esch, M. B. 2021. Critical Considerations for the Design of Multi-Organ Microphysiological Systems (MPS). Frontiers in Cell and Developmental Biology, 9.

March, S., Ramanan, V., Trehan, K., Ng, S., Galstian, A., Gural, N., Scull, M. A., Shlomai, A., Mota, M. M., Fleming, H. E., Khetani, S. R., Rice, C. M. & Bhatia, S. N. 2015. Micropatterned coculture of primary human hepatocytes and supportive cells for the study of hepatotropic pathogens. Nature Protocols, 10, 2027–2053.

NC3RS. 2022. ARRIVE Guidelines [Online]. Available: https://arriveguidelines.org/ [Accessed 08 July 2022].

Olson, H., Betton, G., Robinson, D., Thomas, K., Monro, A., Kolaja, G., Lilly, P., Sanders, J., Sipes, G., Bracken, W., Dorato, M., Van Deun, K., Smith, P., Berger, B. & Heller, A. 2000. Concordance of the toxicity of pharmaceuticals in humans and in animals. Regul Toxicol Pharmacol, 32, 56–67.

Pamies, D., Barreras, P., Block, K., Makri, G., Kumar, A., Wiersma, D., Smirnova, L., Zang, C., Bressler, J., Christian, K. M., Harris, G., Ming, G. L., Berlinicke, C. J., Kyro, K., Song, H., Pardo, C. A., Hartung, T. & Hogberg, H. T. 2017. A human brain microphysiological system derived from induced pluripotent stem cells to study neurological diseases and toxicity. Altex, 34, 362–376.

Pedersen, B. K. & Febbraio, M. A. 2008. Muscle as an endocrine organ: focus on muscle-derived interleukin-6. Physiol Rev, 88, 1379–406.

Peters, M. F., Choy, A. L., Pin, C., Leishman, D. J., Moisan, A., Ewart, L., Guzzie-Peck, P. J., Sura, R., Keller, D. A., Scott, C. W. & Kolaja, K. L. 2020. Developing in vitro assays to transform gastrointestinal safety assessment: potential for microphysiological systems. Lab on a Chip, 20, 1177–1190.

Picollet-D’Hahan, N., Zuchowska, A., Lemeunier, I. & Le Gac, S. 2021. Multiorgan-on-a-Chip: A Systemic Approach To Model and Decipher Inter-Organ Communication. Trends in Biotechnology, 39, 788–810.

Rogozhnikov, D., O’Brien, P. J., Elahipanah, S. & Yousaf, M. N. 2016. Scaffold Free Bio-orthogonal Assembly of 3-Dimensional Cardiac Tissue via Cell Surface Engineering. Scientific Reports, 6, 39806.

Sachs, N., Papaspyropoulos, A., Zomer-Van Ommen, D. D., Heo, I., BÖTtinger, L., Klay, D., Weeber, F., Huelsz-Prince, G., Iakobachvili, N., Amatngalim, G. D., De Ligt, J., Van Hoeck, A., Proost, N., Viveen, M. C., Lyubimova, A., Teeven, L., Derakhshan, S., Korving, J., Begthel, H., Dekkers, J. F., Kumawat, K., Ramos, E., Van Oosterhout, M. F., Offerhaus, G. J., Wiener, D. J., Olimpio, E. P., Dijkstra, K. K., Smit, E. F., Van Der Linden, M., Jaksani, S., Van De Ven, M., Jonkers, J., Rios, A. C., Voest, E. E., Van Moorsel, C. H., Van Der Ent, C. K., Cuppen, E., Van Oudenaarden, A., Coenjaerts, F. E., Meyaard, L., Bont, L. J., Peters, P. J., Tans, S. J., Van Zon, J. S., Boj, S. F., Vries, R. G., Beekman, J. M. & Clevers, H. 2019. Long-term expanding human airway organoids for disease modeling. Embo j, 38.

Sasserath, T., Rumsey, J. W., Mcaleer, C. W., Bridges, L. R., Long, C. J., Elbrecht, D., Schuler, F., Roth, A., Bertinetti-Lapatki, C., Shuler, M. L. & Hickman, J. J. 2020. Differential Monocyte Actuation in a Three-Organ Functional Innate Immune System-on-a-Chip. Advanced Science, 7, 2000323.

Sato, T., Vries, R. G., Snippert, H. J., Van De Wetering, M., Barker, N., Stange, D. E., Van Es, J. H., Abo, A., Kujala, P., Peters, P. J. & Clevers, H. 2009. Single Lgr5 stem cells build crypt-villus structures in vitro without a mesenchymal niche. Nature, 459, 262–265.

Schneider, S., Bubeck, M., Rogal, J., Weener, H. J., Rojas, C., Weiss, M., Heymann, M., Van Der Meer, A. D. & Loskill, P. 2021. Peristaltic on-chip pump for tunable media circulation and whole blood perfusion in PDMS-free organ-on-chip and Organ-Disc systems. Lab on a Chip, 21, 3963–3978.

Shankaran, A., Prasad, K., Chaudhari, S., Brand, A. & Satyamoorthy, K. 2021. Advances in development and application of human organoids. 3 Biotech, 11, 257.

Sharifi, F., Firoozabadi, B. & Firoozbakhsh, K. 2019. Numerical Investigations of Hepatic Spheroids Metabolic Reactions in a Perfusion Bioreactor. Front Bioeng Biotechnol, 7, 221.

Sutherland, M. L., Fabre, K. M. & Tagle, D. A. 2013. The National Institutes of Health Microphysiological Systems Program focuses on a critical challenge in the drug discovery pipeline. Stem Cell Res Ther, 4 Suppl 1, I1.

Takebe, T., Sekine, K., Enomura, M., Koike, H., Kimura, M., Ogaeri, T., Zhang, R.-R., Ueno, Y., Zheng, Y.-W., Koike, N., Aoyama, S., Adachi, Y. & Taniguchi, H. 2013. Vascularized and functional human liver from an iPSC-derived organ bud transplant. Nature, 499, 481–484.

Tamaki, C., Nagayama, T., Hashiba, M., Fujiyoshi, M., Hizue, M., Kodaira, H., Nishida, M., Suzuki, K., Takashima, Y., Ogino, Y., Yasugi, D., Yoneta, Y., Hisada, S., Ohkura, T. & Nakamura, K. 2013. Potentials and limitations of nonclinical safety assessment for predicting clinical adverse drug reactions: correlation analysis of 142 approved drugs in Japan. J Toxicol Sci, 38, 581–98.

Tissuse Gmbh. 2022. Available: https://www.tissuse.com/en/ [Accessed].

Van Norman, G. A. 2019a. Limitations of Animal Studies for Predicting Toxicity in Clinical Trials: Is it Time to Rethink Our Current Approach? JACC: Basic to Translational Science, 4, 845–854.

Van Norman, G. A. 2019b. Phase II Trials in Drug Development and Adaptive Trial Design. JACC Basic Transl Sci, 4, 428–437.

Vernetti, L., Gough, A., Baetz, N., Blutt, S., Broughman, J. R., Brown, J. A., Foulke-Abel, J., Hasan, N., In, J., Kelly, E., Kovbasnjuk, O., Repper, J., Senutovitch, N., Stabb, J., Yeung, C., Zachos, N. C., Donowitz, M., Estes, M., Himmelfarb, J., Truskey, G., Wikswo, J. P. & Taylor, D. L. 2017. Functional Coupling of Human Microphysiology Systems: Intestine, Liver, Kidney Proximal Tubule, Blood-Brain Barrier and Skeletal Muscle. Scientific Reports, 7, 42296.

Ware, B. R., Liu, J. S., Monckton, C. P., Ballinger, K. R. & Khetani, S. R. 2021. Micropatterned Coculture With 3T3-J2 Fibroblasts Enhances Hepatic Functions and Drug Screening Utility of HepaRG Cells. Toxicological Sciences, 181, 90–104.

